# Phosphoproteomics after nitrate treatments reveal an important role for PIN2 phosphorylation in control of root system architecture

**DOI:** 10.1101/2020.06.22.164640

**Authors:** Andrea Vega, Isabel Fredes, José O’Brien, Zhouxin Shen, Krisztina Ötvös, Eva Benkova, Steven P. Briggs, Rodrigo A. Gutiérrez

## Abstract

Nitrate is an important signaling molecule that commands genome-wide gene expression changes that impact metabolism, physiology, plant growth and development. Although gene expression responses to nitrate at the mRNA level have been characterized in great detail, the impact of nitrate signaling at the proteome level has been much less explored. Most signaling pathways involve post-translational modifications of key protein factors and chiefly among these modifications is protein phosphorylation. In an effort to identify new components involved in nitrate responses in plants, we performed analyses of the *Arabidopsis thaliana* root phosphoproteome in response to nitrate treatments via liquid chromatography coupled to tandem mass spectrometry. We identified 268 phosphoproteins that show significant changes at 5 min or 20 min after nitrate treatments. The large majority of these proteins (96%) are coded by genes that are not modulated at the expression level in response to nitrate treatments in publicly available transcriptome data. Proteins identified by 5 min include potential signaling-components such as kinases or transcription factors. In contrast, by 20 min, proteins identified were associated with protein binding, transporter activity or hormone metabolism functions. Interestingly, the phosphorylation profile of *NITRATE TRANSPORTER 1*.*1* (*NRT1*.*1)* mutant plants in response to nitrate at 5 min was significantly different (95%) as compared to wild-type plants. This result is consistent with the role of NRT1.1 as a key component of a nitrate signaling pathway that involves phosphoproteomic changes. Our integrative bioinformatics analysis highlights auxin transport as an important mechanism modulated by nitrate signaling at the post-translational level. We experimentally validated the role of PIN2 phosphorylation in both primary and lateral root growth responses to nitrate. Our data provide new insights into the phosphoproteome and identifies novel protein components that are regulated post-translationally, such as PIN2, in nitrate responses in *Arabidopsis thaliana* roots.

## Introduction

Nitrogen (N) is the mineral nutrient required in the greatest amounts by plants. N is often scarce in natural and agricultural systems, constituting a major factor limiting plant growth and agricultural yield. During the last 50 years, global demand for synthetic N fertilizers has dramatically increased in response to growing agricultural demand. Depending on soil conditions and plant species, less than 50% of the applied N fertilizer is taken up by crops. Excess N may contaminate aquatic systems^1^ or be released into the atmosphere as N-oxide gases^2,3^, both leading to detrimental effects on the environment and human health.

The relevance of N for plants is exemplified by its effects on leaf growth^4^, senescence^5^, root system architecture^6,7^, and flowering time^8,9^. Due to its importance, plants have evolved sophisticated mechanisms to adapt to fluctuating N availability. Furthermore, growth and developmental processes can be regulated by varying the amount of N supplied to plants. For instance, exogenous nitrate applications stimulate lateral root elongation, enabling root growth and colonization in nitrate-rich soil patches^10,11^. However, high nitrate concentrations reduce primary and lateral root elongation under homogeneous growth-conditions^12^. Nitrate is the main form of inorganic N for plants in natural and agricultural soils^13,14^. Besides its nutritional role, nitrate acts as a signaling molecule that regulates several genes involved in a wide range of biological processes^15,16^. With advances in genomic technologies and system approaches, thousands of nitrate-responsive genes have been identified in *Arabidopsis thaliana* roots and shoots^17–24^. These N-response genes include nitrate transporters, nitrate reductase (NR) and nitrite reductase (NiR), putative transcription factors, and stress response genes, as well as genes whose products play roles in glycolysis, N metabolism, and hormone pathways. Moreover, nitrate elicits local and systemic signals to synchronize its availability with plant growth and development^25–29^. Although transcriptional responses activated by nitrate have been described in great detail, it is clear that regulation at the post-translational level is key for N-responses^30,31^.

The role of protein phosphorylation in response to nitrate was initially identified in post-translational modifications in N metabolism. The activity of NR, the enzyme that catalyzes the first step of nitrate reduction, is modulated by protein phosphorylation and then inhibited by 14-3-3 protein interaction^32,33^. Studies in spinach leaves using ^32^P labeling and kinase assays demonstrated that the regulation of NR by light/dark and photosynthetic activity involves protein phosphorylation^34,35^. A subsequent study showed that a 14-3-3 family protein interacts with and inactivates phosphorylated NR in the presence of covalent ions ^32,36^. Earlier experiments also indicated that changes in gene expression in response to nitrate treatments require kinase and phosphatase activities. In maize leaves for example, treatments with inhibitors of calmodulin-dependent protein kinases repress nitrate induction of genes encoding nitrate assimilatory enzymes such as NR, NiR, glutamine synthetase 2 (GS2) and ferredoxin glutamate synthase (Fd-GOGAT)^37^. Conversely, inhibition of protein phosphatases blocked the nitrate-response of NR, NIR and GS2^37^. In another study, pharmacological inhibitors of serine-threonine protein phosphatase and tyrosine protein kinases repressed the nitrate-induced accumulation of transcripts for NR and NiR in barley leaves^38^. These early experiments suggested that changes in the status of protein phosphorylation were important for the regulation of gene expression in response to nitrate treatments.

The discovery that a kinase protein complex can directly phosphorylate the nitrate transceptor NRT1.1/NPF6.3 demonstrated that phosphorylation plays an important role in nitrate signaling. Under low-nitrate conditions, NRT1.1/NPF6.3 is phosphorylated in a threonine residue (T101) by CIPK23-CBL9 complex (CIPK, CLB-interacting protein kinase; CBL, Calcineurin B-like protein), shifting into a high-affinity nitrate transporter^31,39^. In contrast, at high-nitrate levels, NRT1.1/NPF6.3 is dephosphorylated at T101 and turns into a low-affinity transporter. Experiments with a mutant mimicking the phosphorylated form of the transceptor showed the importance of this phosphorylation for the regulation of gene expression at low nitrate concentrations^39^. Phosphorylation of NRT1.1/NPF6.3 also appears to play a role in the modulation of auxin transport and repression of lateral root emergence under low-nitrate conditions^40,41^. Conversely, the dephosphorylated form of NRT1.1/NPF6.3 is critical for up-regulation of *NITRATE TRANSPORTER 2*.*1* (*NRT2*.*1*) gene expression in response to nitrate^39,40^. Bouguyon *et al*. (2015)^40^ studied different mutant alleles of NRT1.1/NPF6.3 (T101D, T101A and P492L substitution), and proposed a different NRT1.1/NPF6.3-dependent signaling mechanism. The short-term induction of *NRT2*.*1* at high nitrate is negatively affected by T101D substitutions but not by P492L and T101A. Long-term regulation of *NRT2*.*1* transcripts at high nitrate and repression of lateral root emergence at low nitrate showed an opposite pattern, where signaling was suppressed by both T101A and P492L mutations but not affected by T101 substitution. Another kinase involved in signaling is CIPK8^42^. In *cipk8* mutants, the rapid induction of genes or primary nitrate response was strongly reduced (40-65% of WT levels), particularly in the low-affinity phase^42^. Both CIPK8 and CIPK23 are rapidly induced by nitrate treatments and downregulated in the *chl1-5* and *chl1-9* mutants, respectively^39,42^. More recently, the calcium sensor CBL1 and the protein phosphatase 2C (ABA-insensitive) ABI2 were also identified as components of this signaling pathway, which regulates NRT1.1/NPF6.3 transport and sensing^43^. The calcium sensor CBL1 also interacts with CIPK23 and this complex was dephosphorylated by ABI2^43^.

In higher plants, CBL/CIPK complexes sense and decode Ca^2+^ signals, triggering specific transduction pathways (reviewed by ^44^). Recent studies have shown that Ca^2+^ plays a role in nitrate signaling transduction and is important for the primary nitrate response in *Arabidopsis* roots^45^. Calcium is a key secondary messenger that triggers changes in signaling pathways, including changes in phosphorylation levels^46,47^. More recently, results published by Liu et al. (2017)^30^ have contributed to our understanding of the relationship between Ca^2+^ signaling and the first layer of transcriptional regulators. They used the luciferase (LUC) reporter gene NIR-LUC, which exhibits a physiological nitrate response in transgenic *Arabidopsis* plants, to identify three CPKs (CPK10, CPK30, and CPK32) that activated the NIR-LUC reporter in an effective and synergistic manner. Additionally, CPK10, CPK30, and CPK32 phosphorylated the transcription factor NIN-LIKE PROTEIN 7 (NLP7) in a Ca^2+^ dependent manner^30^, suggesting that CPK-NLP signaling is a key regulator of primary nitrate responses^48^. All these studies provide evidence that NRT1.1/NPF6.3, calcium and phosphorylation of target proteins are key elements of a signaling pathway involved in the nitrate response.

Global-scale proteomic analysis performed in *Arabidopsis* seedlings, mostly shoot organs, showed that nitrogen starvation and resupply (nitrate or ammonium) modulates protein phosphorylation over a time course of 30 min^49^. In general, proteins such as receptor kinases and transcription factors change their phosphorylation levels after nitrogen resupply at 5-10 min (fast response). Proteins involved in protein synthesis and degradation, central and hormone metabolism showed changes in their phosphorylation level after 10 min (late response). Another study showed that nitrate deprivation affects both protein abundance and phosphorylation status^50^. Nitrate deprivation assays revealed that some proteins, mostly involved in transport, contain sites that are dephosphorylated early in the response ^50^.

In this study, we performed quantitative time-course analyses of the *Arabidopsis* root phosphoproteome in response to nitrate via liquid chromatography coupled to tandem mass spectrometry detection (HPLC-MS/MS). We chose to focus on root-phosphoproteomics profiling in response to nitrate for several reasons: (i) phosphoproteomics and proteomics studies describe phosphorylation levels as more dynamic and mainly independent of protein abundance^51–53^, suggesting that many proteins are regulated by phosphorylation independent of their changes in protein abundance. (ii) Previous global studies of N treatment focused on the proteome and phosphoproteome in *Arabidopsis* seedlings, which interrogates mostly shoot tissues^49,50^. In order to search for new N-regulatory factors, our experimental approach focused on *Arabidopsis* roots because early sensing and responses to N-supply occur in the roots. Several studies have shown that HPLC-MS/MS provides accurate estimates of dynamic phosphorylation levels *in vivo*^54–57^.

We used HPLC-MS/MS to identify phosphorylated proteins with differential profiles in response to nitrate treatments at 0, 5 or 20 min. We found that the nature of these phosphorylated proteins differed significantly from those encoded by genes implicated in nitrate via transcriptomic studies. We found different types of phosphoproteins changing at 5 or 20 min after nitrate treatments. Interestingly, the large majority of these changes depend on NRT1.1/NPF6.3. Kinases and transcription factors were over-represented at 5 min, while proteins involved in protein binding and transporter activity were common by 20 min of nitrate treatments. We found several phosphoproteins involved in auxin transport, including the auxin efflux-carriers PIN2 and PIN4. We validated the role of PIN2 and found dephosphorylation of PIN2 to be important for modulation of root system architecture in response to nitrate. Our analysis reveals that the nitrate signaling pathway mediated by NRT1.1/NPF6.3 leads to important changes in protein phosphorylation patterns and proposes new players that participate in the developmental responses to nitrate in plants.

## Results

### Phosphoproteome analysis of *Arabidopsis* roots in response to nitrate treatments

In an effort to identify new components involved in nitrate responses in plants, we performed large-scale mass spectrometry-based phosphoproteome experiments following nitrate treatments in *Arabidopsis* roots. *A. thaliana (L*.*) Columbia-0* (Col-0) seedlings were grown hydroponically, with a full nutrient solution (Murashige and Skoog basal medium without N) containing 1 mM ammonium as the only N source for 14 days (time 0, see experimental procedures). Two-week-old plants were treated with 5 mM KNO_3_ or KCl, as control. We and other laboratories have used this experimental setup in prior studies because it elicits robust primary nitrate responses in *Arabidopsis* plants^7,16,45,58,59^. A label-free high-performance liquid chromatography-tandem mass spectrometry (HPLC-MS/MS) method was used to identify changes in protein phosphorylation at 0, 5 or 20 minutes after nitrate treatments. We chose these time points because they have shown to be effective in describing transient and persistent protein phosphorylation responses^49,55^. Moreover, this experimental design allows for comparison with transcriptomics data obtained using the same experimental conditions^7,45,58^. For phosphoproteome analysis, we used a previously validated experimental pipeline^52,53,60^. Briefly, phosphopeptides were enriched using cerium oxide affinity capture and analyzed with a HPLC-MS/MS instrument (Figure S1). The spectra were assigned to specific peptide sequences by the MASCOT search engine (FDR < 0.1%). We quantified the relative abundance of each phosphoprotein using average normalized spectral counts (SPCs) of the total number of spectral-peptide matches to protein sequences in three independent biological replicates for each treatment condition. In total, we identified and measured 6,560 unique phosphopeptides which unambiguously mapped to 2,048 phosphoprotein groups (Supplemental Table S1). The majority of identified phosphopeptides (82%) were phosphorylated in a single residue (Figure S2A). The relative distribution of each phosphorylated residue – 80% serine, 18% threonine, and 2% tyrosine (Figure S2B) – was consistent with prior plant phosphoproteomic studies^50,54,61^. The identified phosphopeptides were mapped and grouped in phosphoprotein groups, where proteins that shared peptides were clustered together. A group leader was assigned to each group, based on having the highest number of peptide identifications; throughout the remainder of the article, “phosphoproteins” is synonymous with “group leaders”. The majority of the identified phosphoproteins present one (50%), two (25%) or three (12%) phosphorylated peptides (Figure S2C) with similar distributions of phosphorylated residue (69% Ser, 28% Thr and 3% Tyr; Figure S2D). We recognized phosphoproteins across several biological process, subcellular compartments and cellular functions based on the Gene Ontology (GO) classification (Figure S3). No overrepresented GO categories were observed when comparing against the *Arabidopsis* genome, showing that our experimental strategy was unbiased with regards to annotated protein functions, subcellular locations or biological processes and represents an unbiased *Arabidopsis* proteome sampling.

To identify nitrate-regulated phosphoproteins in *Arabidopsis* roots we performed statistical analysis using analysis of variance (significance: *p* < 0.05). We identified 268 phosphoproteins that were significantly altered under our experimental conditions. We found 62 phosphoproteins regulated at 5 min after nitrate treatments, 40 of which were induced (Figure 1A). We found 152 phosphoproteins differentially regulated at 20 min, 113 of which were induced by the nitrate treatments. The large majority of phosphoproteins were found to be nitrate-regulated at only one time-point, indicating that most changes in the phosphoproteome are transient with an early (5 min) and late (20 min) component in response to nitrate treatments. Two previous studies characterized the phosphoproteome in response to nitrate-depletion^49^ or nitrate-resupply^50^ (Figure 1C). Response to nitrate-resupply resulted in 10 common phosphoproteins and nitrate-deprivation in 6 common proteins. These proteins were mostly associated with nitrate metabolism, such as transporters (NRT2.1; ammonium transporter 1.3, AMT1.3), the *Arabidopsis* H^+^-ATPase 2 (AHA2) proton pump, and nitrate reductase 2 (NIA2). The comparison of the phosphoproteins identified here with transcriptome data revealed that 96% of the phosphoproteins are encoded by genes that do not change expression at the mRNA level (Figure S4). We compared our results with an integrated analysis of available root microarray data under contrasting nitrate conditions (27 experimental datasets corresponding to 131 arrays^62^). This study identified a group of 2286 nitrate-responsive genes regulated at transcription levels. Only 11 of these genes were found in our phosphoproteomic dataset.

**Figure 1.**
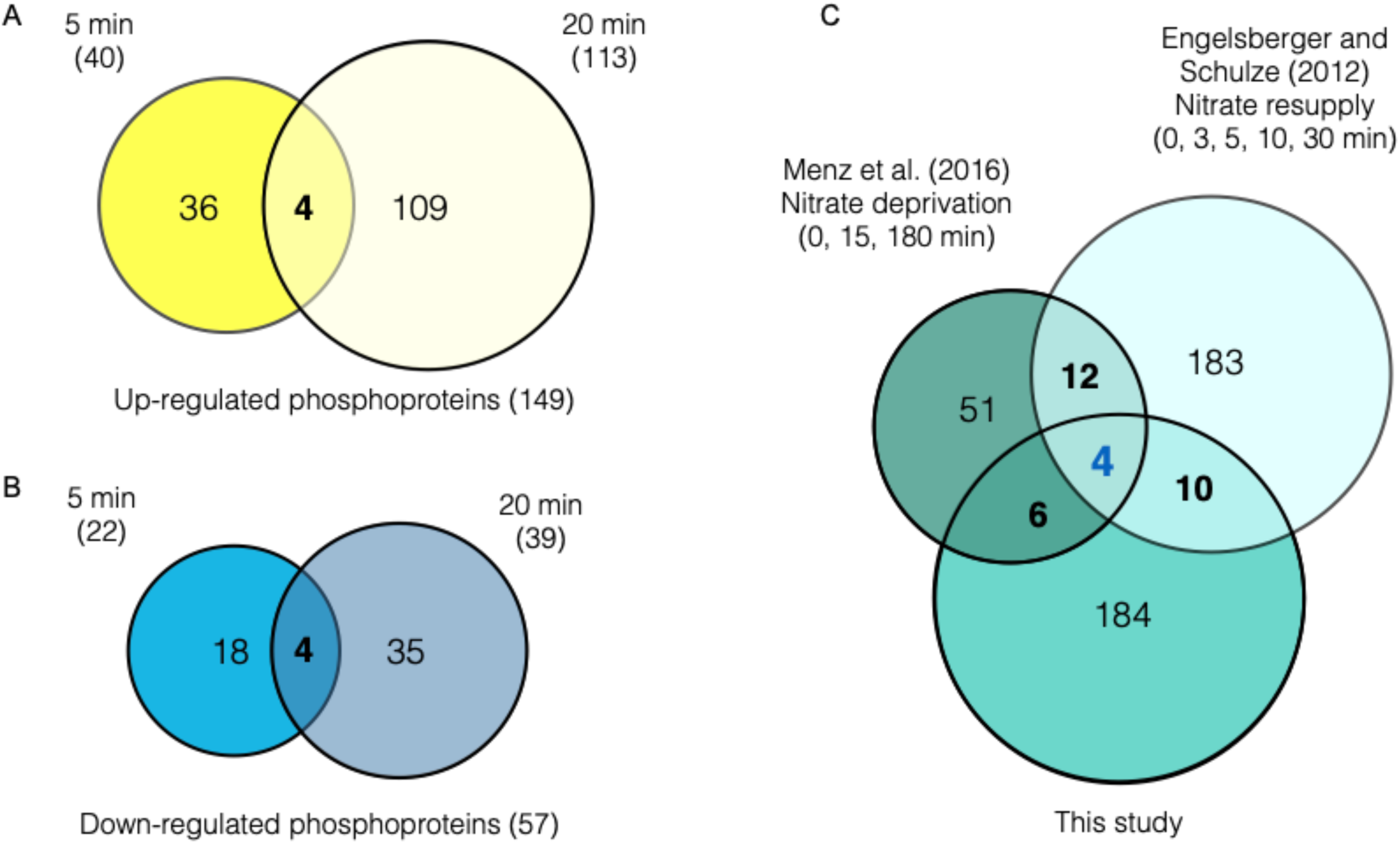
Characterization of the phosphoproteome profile in *Arabidopsis* roots in response to nitrate. Venn diagrams show the overlap of transient (5 min) and more persistent (20 min) changes in phosphoprotein relative-abundance after 5 mM nitrate treatments compared with each control condition (5 mM KCl) in *Arabidopsis* roots. (A) Overlap of phosphoproteins that were “up-regulated” by nitrate at 5 min (dark yellow) and 20 min (light yellow) (B) Overlap of phosphoproteins that were “down-regulated” by nitrate at 5 min (blue) and 20 min (light blue). (C) Identification of common phosphoproteins with published phosphoproteome data-sets in response to nitrate-deprivation^50^ or nitrate-resupply^49^.

Our approach identified new genes coding for phosphoproteins putatively involved in nitrate responses. Changes in phosphorylation patterns improve our understanding of signaling mechanisms that connect nitrate transporters/sensors with transcriptomic response and other biological processes. The small overlap between the phosphoproteomic studies to date highlights the importance of ours and additional future studies to address this important aspect of posttranslational modifications in response to N-signals in plants.

### Functional enrichment in the phosphoproteome reveals distinctive signaling and regulatory processes occurring in early and late responses to nitrate

To evaluate the biological significance of the phosphoproteome patterns observed in response to nitrate, hierarchical clustering analysis was performed on the phosphoprotein dataset at 5 and 20 min following nitrate treatments in Col-0 roots (Figure 2). In order to identify the most prominent functional categories affected, we searched for overrepresented biological terms in each cluster using the BioMaps program^63^ and the PANTHER classification system^64,65^ (significance: *p* < 0.05, corrected by FDR). This analysis highlighted several signaling, regulatory or metabolic functions differentially associated with early (5 min) and late (20 min) components in response to nitrate treatments. “DNA binding” and “Nucleic acid binding” categories were overrepresented in cluster 6, containing phosphoproteins that increase their levels at 5 min and do not change at 20 min. This cluster also includes previously undescribed transcription factors (TFs) in the N-response cascade in diverse transcriptomic studies^21–24,66^. In contrast, cluster groups containing mostly phosphoproteins that changed their levels at 20 min in response to nitrate were enriched in the functional categories: “transport” and “nitrogen compounds metabolism” (clusters 2 and 8). Consistent with this, increases in phosphoprotein levels in response to nitrate were observed in NRT2.1, the lysine–histidine-like transporter 4 (LHT4), the UDP-Arabinofuranose Transporter 4, AMT1-3 and the K+-transporter 8 (KUP8) only at 20 min following nitrate treatment. Increased abundance of the phospho-peptide Ser-501, corresponding to NRT2.1, was observed at 20 min after nitrate treatment. This site is localized in a C-terminal phosphorylation hotspot (Phosphorylation site database and predictor Phosphat4.0^67,68^) and was also identified in nitrate-deprivation experiments with an opposite phosphorylation response^50^. Also, phosphopeptides for AMT1.3 with phosphorylation at Thr-464 and Ser-487 were identified as “up-regulated” by nitrate treatments at 20 min. The phosphorylation of both sites inhibits transport function^69^ and were also identified as phosphopeptides in nitrate deprivation^50^ and resupply^49^ experiments. We also found that nitrate strongly increased levels of phosphorylated nitrate reductase NIA2 at the highly conserved and regulatory site Ser-534^70^. These results are consistent with previous studies and suggest overall regulation of N metabolism by phosphorylation of key players by 20 min after nitrate treatments.

**Figure 2.**
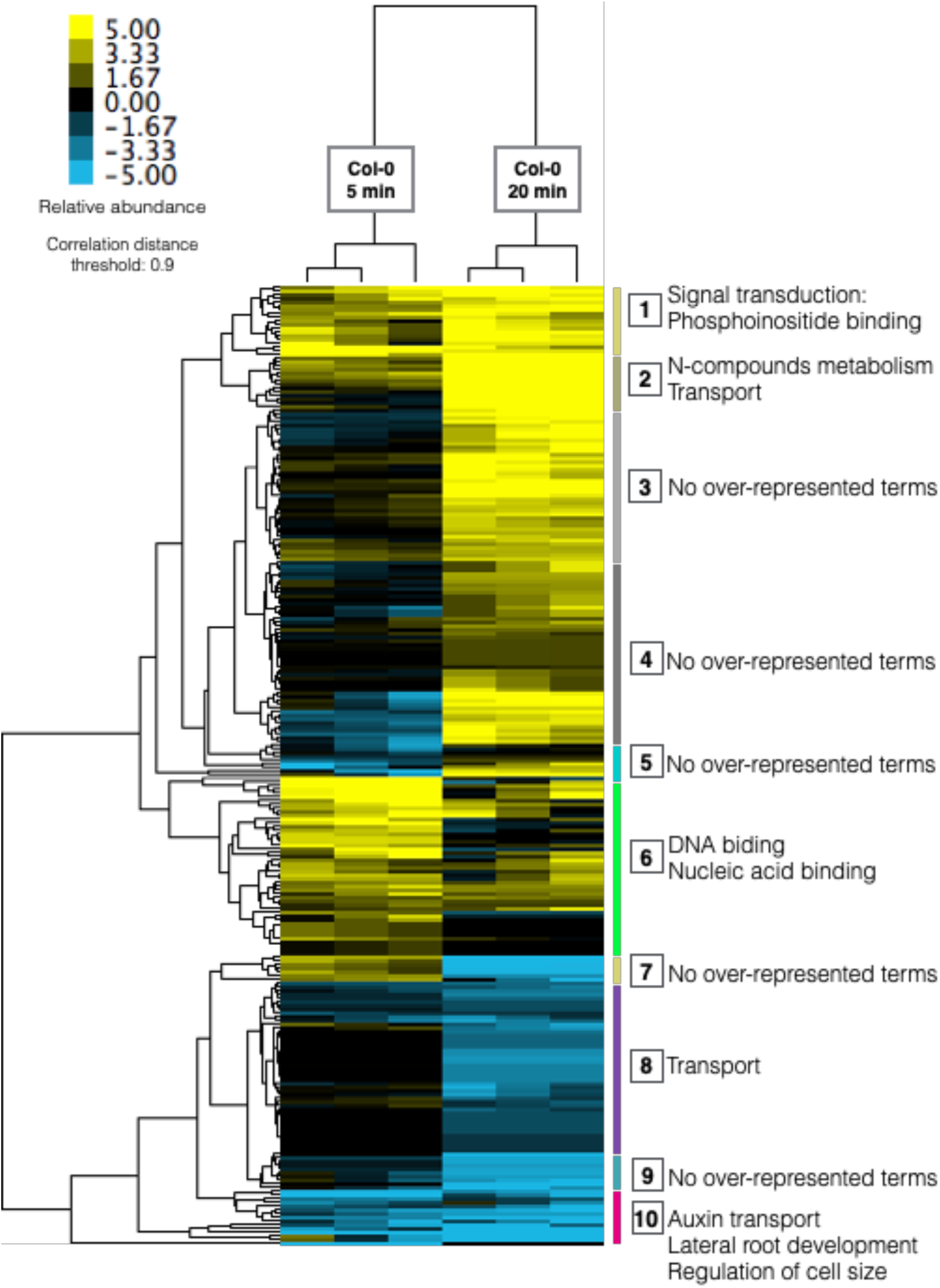
Functional analysis of the phosphoproteome profile reveals signaling and regulatory processes occurring in early and late response to nitrate. The Figure showed a hierarchical clustering of nitrate-phosphoproteins with differential abundance at 5- or 20-min in response to nitrate. Phosphoproteins were normalized by Z-score and clustered using the Euclidean distance method with average linkage. The resulting clusters are shown as a heat map, where vertical bars and numbers to the right of the map denote the group (composed of all terminal nodes in the hierarchical tree) of phosphoprotein (correlation > 0.9) with a selected profile in early (5-min) or late (20-min) nitrate-response. Functional Gene Ontology (GO) categories significantly enriched (hypergeometric test with FDR, *p* < 0.05) in each cluster are highlighted to the right of the group.

The final group of clusters showed enrichment in signaling and regulatory pathways, with different profiles at 5 min and 20 min. Clusters 1, 5, and 7, contained phosphoproteins with opposite regulation at 5 and 20 min, and showed proteins related to microRNA processing, phosphoinositide and phosphatidylinositol binding functions. Phosphoinositides can act in signaling pathways and serve as precursors for phospholipase C (PLC)-mediated signaling. A previous study in our laboratory implicated a PLC-activity in the nitrate signaling pathway^45^. Our results are consistent with these results and show that PLC2, represented by the phosphopeptide Ser-280, was identified as up-regulated in response to nitrate at 20 min (in Cluster 1). Cluster 10 is interesting for nitrate responses because it includes many components involved in classical processes regulated by nitrate in *Arabidopsis* roots. For instance, auxin and lateral root development are biological functions enriched in this group, which is consistent with auxin pathways being modulated by nitrate^16,71^. Several reports indicate that auxin acts as regulator of root system architecture in response to nitrate availability^7,71,72^.

Overall, our dataset captures the dynamic effects of N-signaling on phosphoproteome profiles, which implicate a cascade of nonoverlapping processes in early and late responses. The earliest steps in the N phospho-dynamics were involved in signal transduction and transcription factor activity. In contrast, the later phosphoprotein data-set was enriched in metabolic, transport and root developmental processes. These temporal mechanisms show a transition of phosphorylation dynamics from phosphoproteins essential to signaling networks to proteins associated with biological processes involved in nitrate response.

### NRT1.1/AtNPF6.3 is essential for transient phosphoprotein changes in response to nitrate treatments

The only nitrate sensor described to date is the nitrate transporter, NRT1.1/NPF6.3^39,73^. In order to understand the importance of NRT1.1/NPF6.3 for nitrate elicited changes in the phosphoproteome observed, we analyzed the phosphoproteomic profile of roots treated with 5 mM KNO3 or KCl (control) in a *nrt1*.*1-null* background (mutant *chl1-5*), using the same experimental conditions described above. 74% of phosphoproteins were detected in both datasets, yet only 5% of the nitrate-phosphoproteome response observed in wild-type plants was maintained in the *chl1-5* mutant (Figure 3). Moreover, 95% of phosphoprotein levels were altered in the *chl1-5* mutant plants. This result indicates that this gene is important for modulating protein phosphorylation in response to nitrate, in addition its established role as nitrate transceptor.

**Figure 3.**
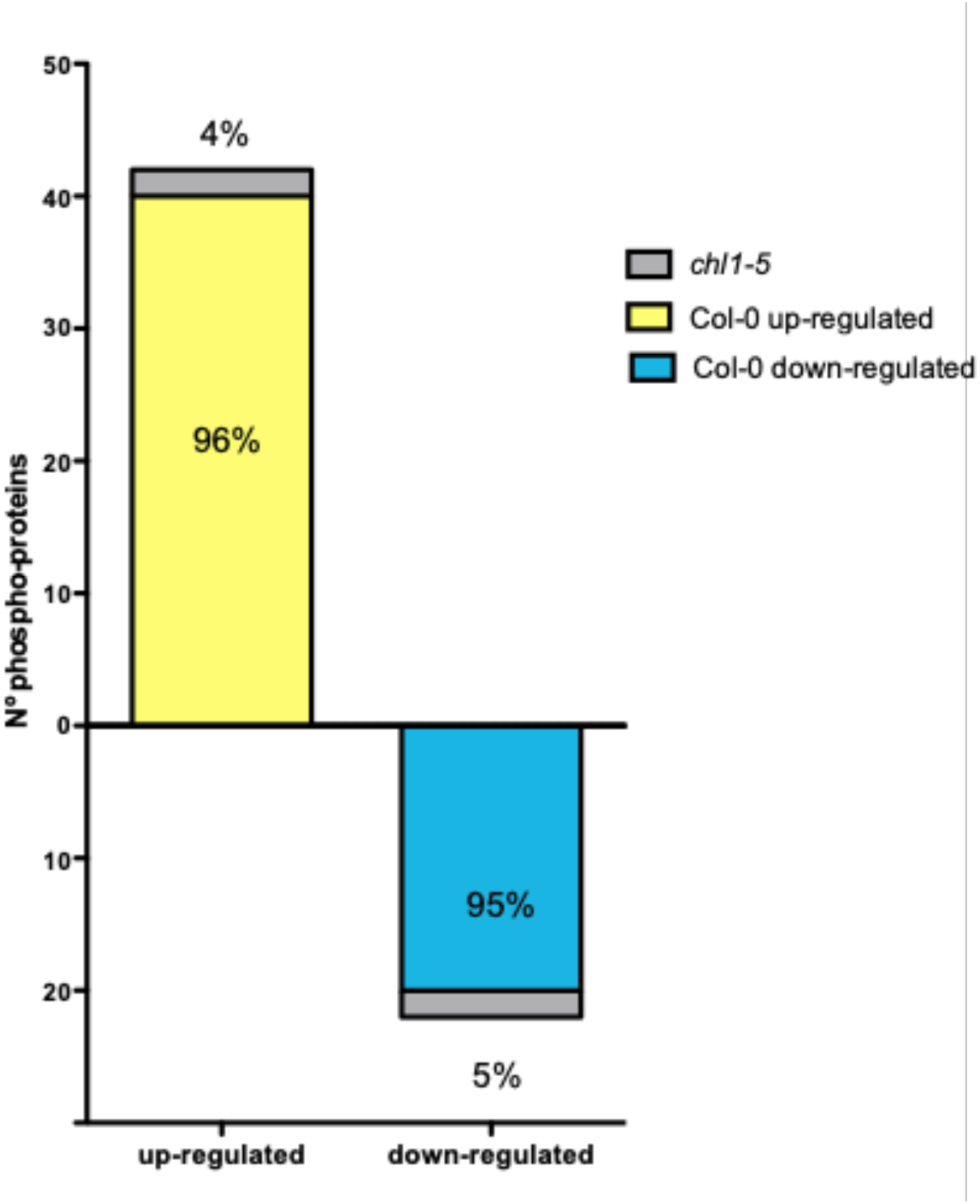
Characterization of the phosphoproteome profile in response to nitrate in roots of *Arabidopsis* (Col-0) or *chl1-5* mutant plants. Percentage of nitrate-phosphoproteome responses at 5 min in wild-type plants that were maintained in *chl1-5*-mutant plant roots.

Our analysis indicates that NRT1.1/NPF6.3 is critical for maintaining the *Arabidopsis* phosphoproteome in response to nitrate availability. It also denotes that NRT1.1/NPF6.3 function is required for rapid changes in phosphorylation of key proteins in response to nitrate in *Arabidopsis* roots. For example, phosphorylated peptides that map to proteins associated with nitrogen metabolism (NRT2.1, AMT1-3 and NIA2) were identified in *chl1-5* mutant roots, but their levels were not affected in response to nitrate.

### Network analysis reveals regulatory sub-networks connected to transcription factors and potential kinases in response to nitrate

To uncover key biological processes modulated by changes in phosphorylation, we performed a multinetwork analysis with our phosphoproteomics data. We generated this network by integrating different levels of information, including protein-protein interactions from BioGRID^74^, predicted protein-DNA interactions of *Arabidopsis* TFs (DapSeq)^75,76^, *Arabidopsis* metabolic pathways (KEGG), and miRNA-RNA, as described previously^16^. We also integrated kinase-substrate predictions and identified the most significant phosphorylation motifs and their predicted kinase families from our phosphoproteomic datasets using the Motif-X algorithm^77^ and the PhosPhAt Kinase-Target interactions database^78^ (Figure S5). We used the Cytoscape^79^ software to visualize the resulting network, wherein genes that encoded each phosphoprotein were represented as nodes linked by edges that signify any of the functional relationships annotated in the various databases indicated above. We generated a network of 206 nodes with 700 interactions (Figure 4A). Although the majority of these genes are not regulated by nitrate at the mRNA level, they form a highly interconnected network which includes potential regulatory transcription factors and kinase components. They are connected by multiple edges, including protein-protein, protein-DNA, and metabolic interactions. This result suggests that the products of these genes form connected biological modules that are coordinately regulated at the post-translational level. This network included several TFs with a high number of regulatory links. The most connected were the Trihelix transcription factor 1 (GTL1), the WRKY DNA-binding protein 65 (WRKY65), and the RELATED TO VERNALIZATION 1 (RTV1) transcription factors, which had not previously been characterized in the context of nitrate response. Intriguingly, GTL1 regulates root hair growth in *Arabidopsis*^80^, which has recently been described as a biological process modulated by nitrate treatments under the same experimental conditions^81^. A previous study indicated that WRKY65 interacts at the protein level with the mitogen-activated protein kinase 10 (MPK10), which binds with the lateral organ boundaries domain 16 (LBD16), LBD18, and LBD29 transcription factors^82,83^. These LBDs are inducible by auxin and play a role in the formation of lateral roots^83^. In addition, MPK10 interacted with other genes involved in the auxin response^82^, while LBD29 regulated genes involved in auxin transport, including auxin efflux-carriers PIN1 and PIN2^83^. This evidence suggests that WRKY65 could be involved in nitrate–auxin signaling crosstalk. These three TFs appear to coordinate different subnetworks largely involved in auxin transport and nitrogen metabolism. Consistent with this observation, analysis of over-represented gene ontology annotations highlights the importance of auxin transport (Figure 4B). Other over-represented biological functions in our network were mRNA binding and splicing, regulation of translation and kinase activity (Figure 4B).

**Figure 4.**
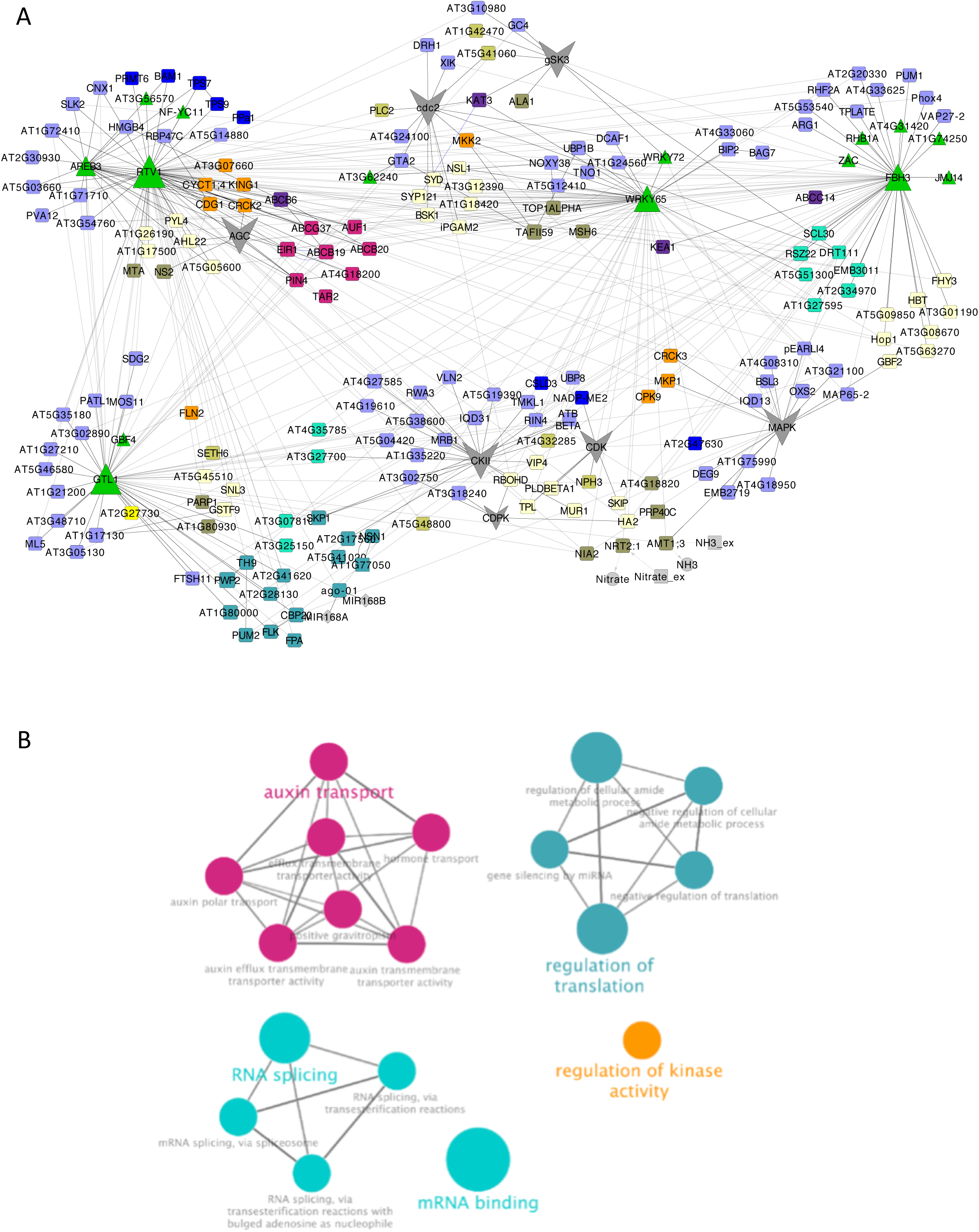
Gene network of nitrate-modulated phosphoproteins. (A) A network for the phosphoproteins (encoded genes) and putative kinases family were represented as nodes with color and shapes assigned according to function (e.g., blue squares: metabolic encoded-genes, green triangle: transcription factors or grey hexagon: kinase families). Edges connecting nodes represent functional interactions: transcription factor/TARGET regulation or predicted kinase-substrate control. The size of the triangle or hexagon is proportional to the number of targets of the TF or putative kinase family, respectively. (B) Over-represented biological processes in the phosphoproteins gene-network in response to nitrate (hypergeometric test with FDR, *p* < 0.05). The network was analyzed using BINGO and ClueGO tools in Cytoscape software. Over-represented gene ontology terms are shown as a node connected by edges based on semantic relationships as defined in the ontology.

Overall, this network analysis highlights a potential role of multiples TFs in linking the N signal and regulatory nitrate responses that show significant enrichment for key functions in signaling pathways and validates the important role auxin plays in the nitrate-response of root system architecture.

### PIN2 is important for modulation of root system architecture in response to nitrate treatments

Auxin is a key phytohormone in plants, involved in growth and developmental responses. Several reports have shown that auxin mediates root developmental responses to nitrate availability^7,16,71,72,84–86^. Nitrate can regulate auxin biosynthesis, transport and accumulation. In response to nitrate, several auxin-related modules are regulated, including the upregulation of auxin receptor AUXIN SIGNALING F-BOX 3 (AFB3) and the feedback regulation by miR393^7,71^. This auxin-signaling component in response to nitrate is implicated in both primary and lateral root growth^7^. Other important evidence comes from the analyses of the *Arabidopsis* transcriptomic response upon nitrate treatments. They show that several genes involved in auxin transport are affected, including auxin efflux-carriers PIN1, PIN4 and PIN7^16,71^. Moreover, the nitrate transceptor NRT1.1/NPF6.3 not only senses and transports nitrate but can also transports auxin, a process that regulates auxin-localization patterns and lateral-root elongation^84^. Consistent with these prior observations, auxin transport was conspicuous throughout our entire phosphoproteomic analysis (Figure 2 and 4). PIN phosphorylation has been shown to be essential for auxin transport and distribution^87–92^. PINs phosphorylation in conserved serine and/or threonine of the central loop controls intracellular trafficking, recycling and polar membrane localization of PIN proteins (Review by^87^). Previous studies indicate that PIN polar localization explains auxin fluxes and distribution patterns, which could mediate differential growth in diverse plant tissue such as roots^91,93^. To validate the relevance of phosphorylation in auxin transport and its connection with nitrate response, we chose the auxin efflux-carrier PIN2 (identified in our experimental dataset) due to its potential role in linking nitrate and changes in root system architecture (RSA). We found an uncharacterized PIN2 phosphorylation site (Ser439) at the end of the hydrophilic cytoplasmic loop (C-loop, Figure 5A). Its phosphopeptide levels decreased by close to 75% in response to nitrate treatments by 5 min (Figure 5B). PIN2 belongs to the PIN-FORMED protein family of auxin transporters and is the principal component mediating basipetal auxin transport in roots^94,95^. This polar auxin transport is essential for root gravitropism^95^ and lateral root formation^96^. Intriguingly, we detected only one phosphorylated peptide for PIN2 in response to nitrate. Protein sequence alignment indicated that this phosphosite is highly conserved in different plant species representing gymnosperm, mono- and dicotyledonous plant lineages of seed plants (Figure 5A). This phosphopeptide has also been described as down-regulated after auxin treatment but its function remains an open question^57^.

**Figure 5.**
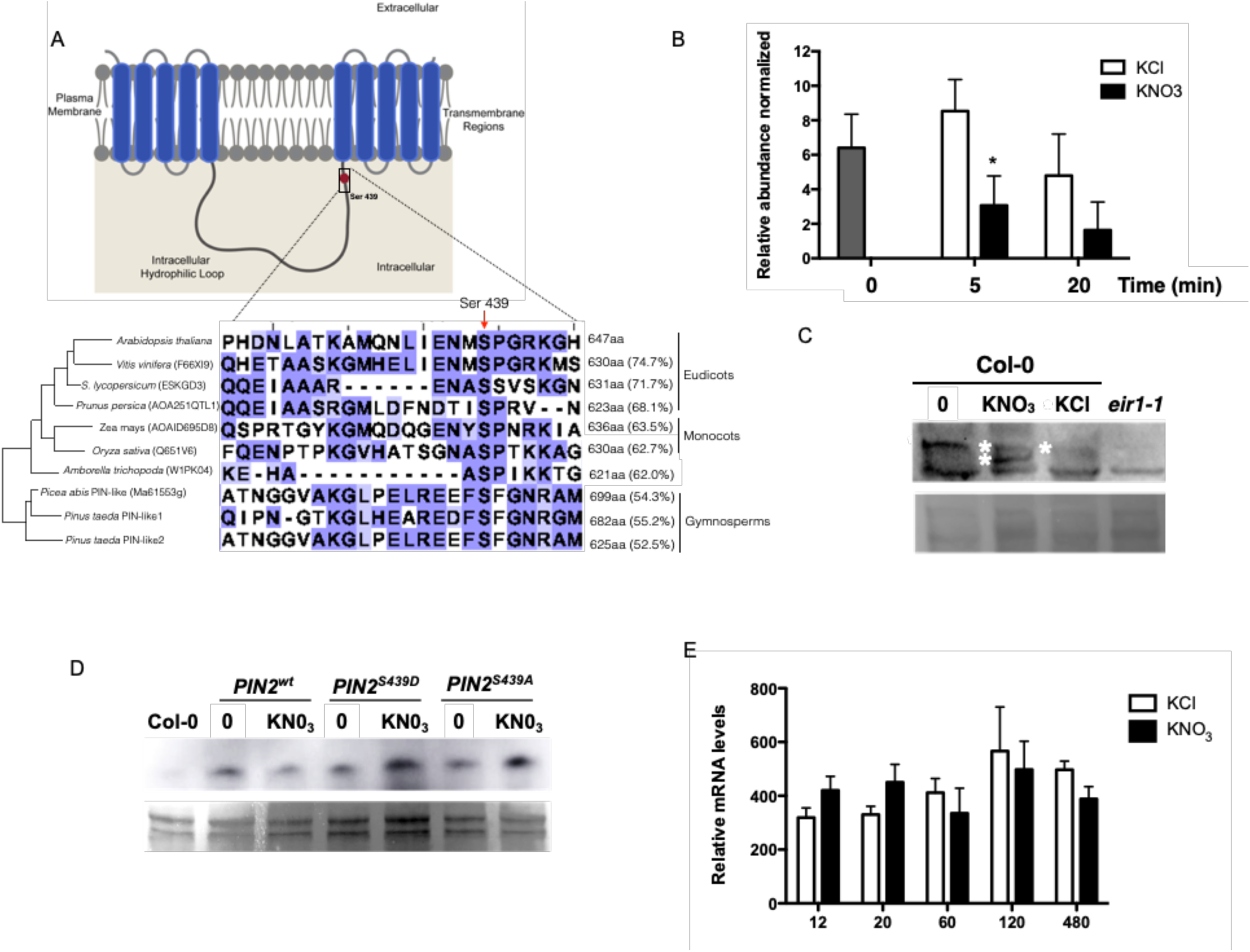
Nitrate regulates PIN2 phosphorylation levels. (A) Schematic representation of the phosphosite (Ser 439) identified in PIN2 in our study. PIN2 protein possesses three specific regions: a region with five transmembrane segments (residues 1–163), a central hydrophilic loop extending from residue 164 to 482 and a region with five additional transmembrane segments (residues 483-647). Protein alignment with PIN2 or PIN-like proteins from different species indicates conservation of the serine residue in eudicot, monocot, and gymnosperm species. The red arrow highlights the S439 in the central loop of PIN2 proteins in different plants. The size of each protein in amino acids and the percentage of identity against the *Arabidopsis* PIN2 are indicated. (B) Levels of PIN2 phosphopeptide in our experiments. Bars represent the mean plus standard error of replicates. The asterisk indicates statistically significant differences (*p* < 0.05). (C) Detection of phosphorylation PIN2 by Phos-tag Western blotting. *Arabidopsis* plants (Col-0) were growth in ammonium as only nitrogen source, and treated with 5 mM KNO3 or 5mM KCl as control. Total protein from roots were analyzed in SDS PAGE using Phos-tag to detect changes in phosphorylation status. Immunoblotting was performed with PIN2 antibody. Total proteins isolated from *eir1*.*1* roots were used as a negative control. (D) Western blot against PIN2 protein comparing nitrate treated (KNO3) and control (0) condition in *Arabidopsis* roots for all genotypes (*eir1-1* mutant background was complemented with PIN2::PIN2^wt^-GFP (PIN2^wt^), PIN2::PIN2^S439D^-GFP (PIN2^S439D^ phospho-mimic point mutation) or PIN2::PIN2^S439A^-GFP (PIN2^S439A^, phospho-null point mutation). (E) Time-course analysis of PIN2 mRNA levels in response to nitrate treatments in *Arabidopsis* roots.

As a first step to understand the function of PIN2 phosphorylation in the nitrate response, we performed a Phos-tag Western Blot analysis to confirm the changes in PIN2 phosphorylation after nitrate treatment. We detected two, fast- and slow-mobility, PIN2 specific bands indicating the presence of two phospho-populations in response to nitrate: one more and another less phosphorylated PIN2. In contrast, only one slow-mobility band corresponding to the more phosphorylated PIN2 subpopulation could be observed at time 0 (ammonium-supplied roots) or under control conditions (KCl-treated roots) (Figure 5C). To assess whether these changes in PIN2 phosphorylation status are a result of a change in protein abundance, we analyzed PIN2-GPF protein levels by Western Blot under our experimental conditions. We introgressed the construct *PIN2::PIN2-GFP* (*PIN2*^*wt*^*-GFP*) into the *pin2* loss-of-function mutant plant *eir1-1*^97^. No differences in protein levels were observed in roots treated with nitrate as compared to roots at time 0 (Figure 5D). To understand the function of this specific PIN2 phosphosite Ser439, we also analyzed the protein levels in *pin2* null mutant complemented with phospho-null (*PIN2::PIN2*^*S439A*^*-GFP*) or phospho-mimic (*PIN2::PIN2*^*S439D*^*-GFP*) versions of PIN2-GFP. Similarly, PIN2 protein concentrations were similar when our experimental conditions mimicked PIN2 phospho-modifications (Figure 5D). Moreover, no regulation at the mRNA level was observed in *PIN2* during nitrate responses (Figure 5E). These results demonstrated that nitrate regulates PIN2 at the posttranslational level, causing de-phosphorylation of PIN2 at a specific phosphosite.

Next, we evaluated the role of PIN2 in root system architecture (RSA) in response to nitrate treatments. We grew wild-type (Col-0) and *pin2* mutant (*eir1*.*1)* plants for 2 weeks on ammonium as sole N source (time 0) and evaluated RSA after nitrate treatments. We measured primary root length 3 days after 5 mM KNO_3_ or KCl treatments. As expected for this experimental set up, we found that nitrate-treated wild-type plants developed shorter primary roots as compared to KCl-treated plants, consistent with earlier results indicating that nitrate treatments inhibit primary root elongation under these experimental conditions^7^ (Figure 6A). However, primary roots of *eir1-1* plants were not significantly inhibited by nitrate treatments as compared to wild-type plants. We also analyzed the density of lateral roots in response to nitrate treatments. In wild-type plants, nitrate treatments increased the number of lateral roots (emerged and initiating) as compared with KCl treatments (Figure 6B). In contrast, the lateral root response to nitrate treatment was altered in the *eir1-1* mutant and the density of lateral roots was significantly reduced as compared with wild-type plants (Figure 6B). These results show that PIN2 plays an important role in modulating RSA in response to nitrate treatments.

**Figure 6.**
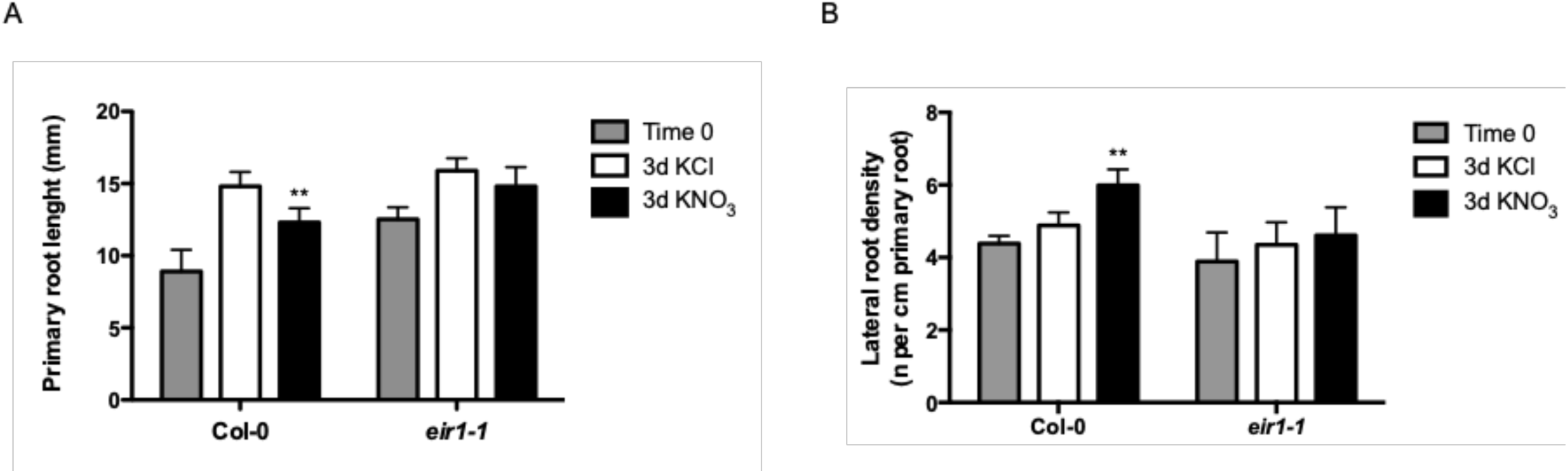
PIN2 is essential for nitrate regulation of primary root growth and lateral root density. (A) Primary root length of Col-0 wild type plants or *eir1-1* mutant plants was measured using the ImageJ program 3 days after 5 mM KNO3 or KCl treatments. (B) Number of initiating and emerging lateral roots of Col-0 or *eir1-1* mutant plants were counted using light microscopy 3 days after 5 mM KNO3 or KCl treatments. Bars represent mean plus standard error of replicate experiments. Asterisk indicates statistically significant difference between means (** *p* < 0.01).

### PIN2 de-phosphorylation of Ser439 plays a role in modulating the root system architecture in response to nitrate treatments

To understand the function of the PIN2 phosphosite identified in this study, we analyzed the *pin2* null mutant *eir1-1* complemented with phospho-null (S439A) or phospho-mimic (S439D) versions of PIN2-GFP. As PIN2 plays an important role in root gravitropism, we evaluated the gravitropic curvature in wild-type, *eir1-1* mutant plants and PIN2-GFP transgenic plants. All seedlings were germinated and grown in MS media in the absence of nitrate for 7 days, and then they were transferred to agar plates with or without nitrate. Agar plates were rotated 90° and root curvature was measured at 24 hours. As expected, *pin2* mutant plants showed an agravitropic phenotype. Both *PIN2*^*S439D*^*-GFP* and *PIN2*^*S439A*^*-GFP* were able to rescue the agravitropic phenotype of *eir1-1* mutant plants. This result indicates that the phosphorylation status of PIN2 at S439 is not relevant for root gravitropic responses in *Arabidopsis* (Figure S6). This result also indicates that phosphorylation at S439 does not impair all PIN2 functions.

Next, we evaluated the RSA in response to nitrate treatments in the same phospho-mimicking (*PIN2*^*S439D*^*-GFP*) or -null (*PIN2*^*S439A*^*-GFP*) genotypes and compared it to wild-type plants. Plants were grown for 2 weeks on ammonium as sole N source and then treated with 5 mM KNO_3_ or KCl for three days. Complementation of the *eir1-1* mutant with a wild-type version of PIN2 (*PIN2*^*wt*^) restored normal RSA responses to nitrate (Figure 7). *eir1-1* mutant plants complemented with *PIN2*^*S439A*^ showed RSA changes similar to wild-type plants in response to nitrate treatments, albeit slightly less pronounced. Three days after nitrate treatments, primary root growth was inhibited 42% by nitrate in the *PIN2*^*S439A*^*-GFP* genotype as compared with nitrate-treated wild-type plants (Figure 7A). Similarly, lateral root density in *PIN2*^*S439A*^*-GFP* plants increased 47% as compared to wild-type plants in response to nitrate treatments (Figure 7B). In contrast at the end of the 3-day treatment, the primary root length and lateral root density did not differ between nitrate or KCl treatment in *eir1-1* mutant plants complemented with *PIN2*^*S439D*^*-GFP*. This result, comparable to the response in *eir1-1* roots, indicates that regulation of PIN2 phosphorylation status at S439 is necessary for normal RSA modulation in response to nitrate treatments.

**Figure 7.**
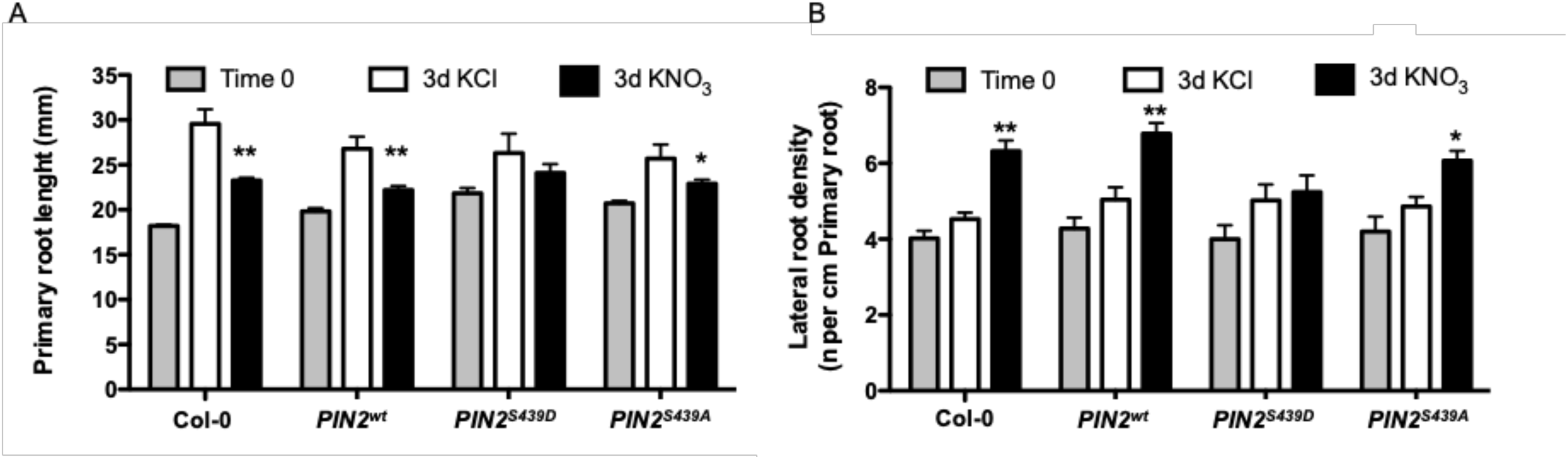
PIN2 dephosphorylation at S439 is important for modulation of primary root growth and lateral root density in response to nitrate treatments. Four different genotypes were used in these experiments: *Arabidopsis* Col-0, *eir1-1* mutant background complemented with PIN2::PIN2^wt^-GFP (PIN2^wt^), PIN2::PIN2^S439D^-GFP (PIN2^S439D^ phospho-mimic point mutation) or PIN2::PIN2^S439A^-GFP (PIN2^S439A^, phospho-null point mutation). All genotypes were grown hydroponically as described in Methods and treated with 5 mM nitrate or KCl (A) Primary root length of the different genotypes was measured using the ImageJ program 3 days after 5 mM KNO3 or KCl treatments (B) Number of initiating and emerging lateral roots for all genotypes were counted using light microscopy 3 days after 5 mM KNO3 or KCl treatments. Bars represent mean plus standard errors. Asterisk indicate statistically significant difference between means (\**p* < 0.05, \*\**p* < 0.01).

### PIN2 phosphosite regulates polar plasma membrane localization in response to nitrate

To explore the impact of nitrate-regulated phosphorylation of PIN2 on cellular localization, we examined the subcellular localization pattern in PIN2-GFP genotypes with phosphosite substitutions. PIN2^WT^-GFP proteins were accumulated at the plasma membrane of epidermal and cortical cells, as previously described (Figure 8A)^86^. PIN2^WT^-GFP fluorescence signal increased in epidermal and cortical cells 2 hours after nitrate treatments as compared to roots in control conditions (roots without nitrate treatments, Figure 8A). Interestingly, mutations at S439 altered this pattern. PIN2^S439A^-GFP plants showed higher fluorescence in the plasma membrane even without nitrate treatment as compared to PIN2^wt^-GFP or PIN2^S439D^-GFP (Figure 8A). The total fluorescence in all experimental conditions analyzed here was similar (Figure 8B). In response to nitrate, PIN2^WT^-GFP and PIN2^S439A^-GFP were accumulated at the plasma membrane of epidermal and cortical cells at comparable levels (Figure 8C and 8D). On the contrary, PIN2^S439D^-GFP plants showed lower levels at epidermal and cortical cells in response to nitrate treatments as compared to PIN2^WT^-GFP or PIN2^S439A^-GFP (Figure 8C and 8D).

**Figure 8.**
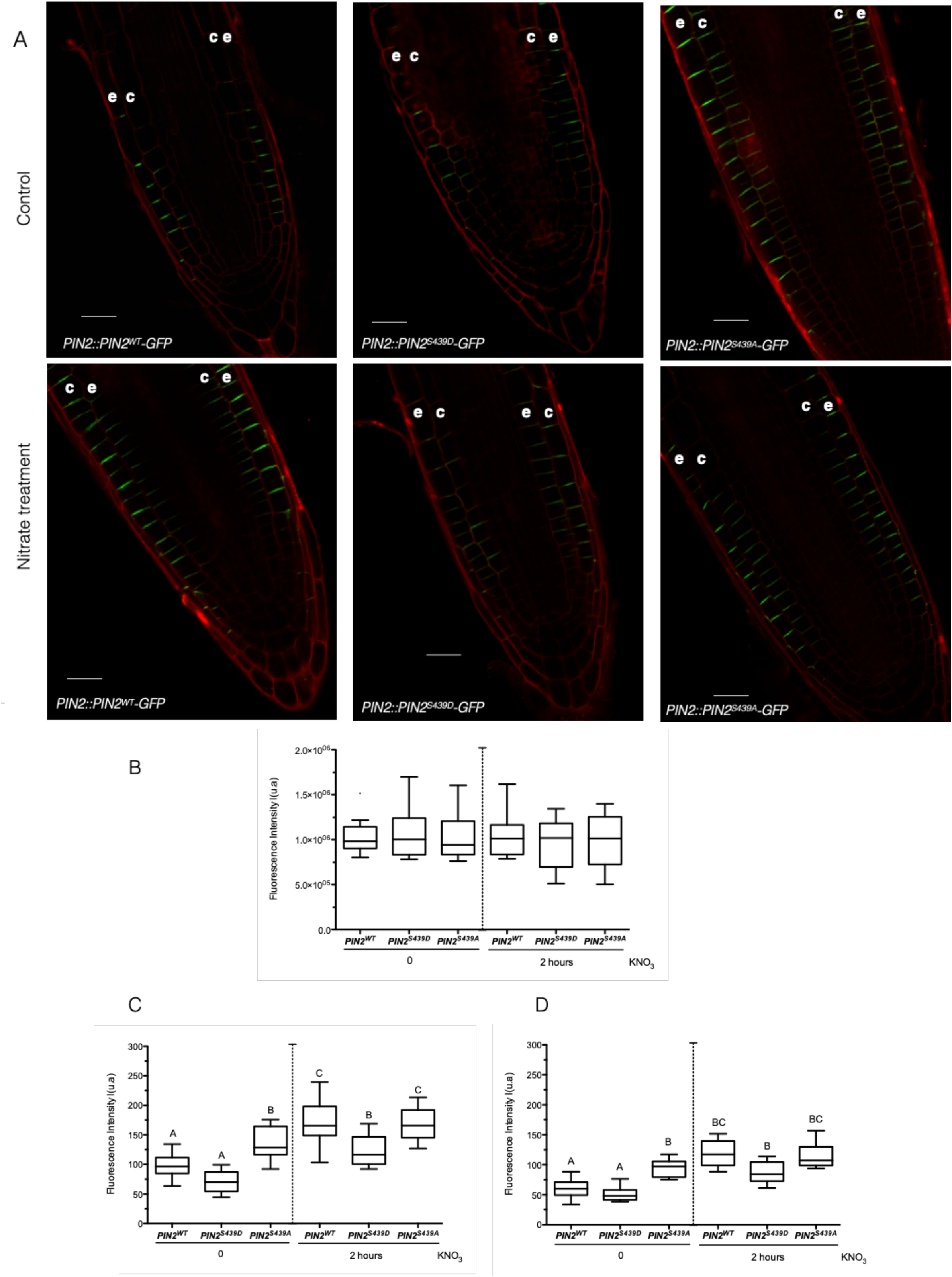
PIN2 phosphorylation at S439 is important for modulation of protein localization in response to nitrate treatments. Three different genotypes were used in these experiments. *eir1-1* mutant background complemented with PIN2::PIN2^wt^-GFP (PIN2^wt^), PIN2::PIN2^S439D^-GFP (PIN2^S439D^ phospho-mimic point mutation) or PIN2::PIN2^S439A^-GFP (PIN2^S439A^, phospho-null point mutation). All genotypes were grown hydroponically as described in Methods and treated with 5 mM nitrate for 2 hours. (A) Confocal microscopy using Zeiss Airyscan microscope and Zeiss-blue3.1 software was used to visualize PIN2 in *Arabidopsis* roots. “e” denotes epidermis and “c” cortex. Scale bar = 20 µm. Tuckey box plots show PIN2-GFP fluorescence intensity quantification (number of roots analyzed per experimental conditions = 8-10; arbitrary units, a.u.) at the total cell membrane in roots (B), and from epidermal (C) and cortex (D) plasma membrane in the different genotypes. Letters denote statistically significant difference between means as determined by ANOVA analysis (*p* < 0.05).

These results indicate that PIN2 phosphorylation status at S439 is important for a correct subcellular localization-pattern in response to nitrate treatments. Moreover, these results indicate that post-translational control impinging upon PIN2 localization is required for RSA changes in response to nitrate treatments.

## Discussion

A key plant nutrient, N also acts as a signal that regulates a myriad of plant growth and developmental processes. Nitrate, a main N-source in natural and agriculture soils, elicits genome-wide changes in gene expression for thousands of genes involved in various biological functions. Nitrate responses have been characterized in great detail at the transcriptome level. However, post-translational modifications have not been characterized in detail. In this study, we evaluated phosphoproteomic profiles in *Arabidopsis* roots in response to nitrate treatments. We focused on characterizing early nitrate-elicited changes in protein phosphorylation. Protein phosphorylation and dephosphorylation plays a central role in modulating protein function in plant signaling-pathways involved in a wide range of processes relating to hormones, nutrients, and responses to stress ^54–57,61,98^.

Our analysis demonstrated that early and late changes in phosphoprotein levels occur in response to nitrate in roots. Furthermore, we identified candidates for nitrate-signaling and biological functions underlying the nitrate response in roots. Interestingly, the majority of proteins and corresponding genes identified in our analysis have not been previously associated with nitrate responses. The phosphoproteomic profile was characteristic of each time-point, showing dynamic changes in phosphorylation patterns in response to nitrate treatments. Early changes in phosphorylation levels (5 min) mainly affected proteins associated with gene regulation, including transcription factors and splicing process. These results are consistent with rapid and dynamic N-responses observed at the mRNA level described in previous studies^21–23,71^. Recent studies have identified new transcription factors^21–23^ involved in the control of gene expression by nitrate. Interestingly, the majority of the transcription factors that we detected as differentially phosphorylated had not been identified as part of the nitrate response. The phosphoproteomic response to nitrate at 20 min revealed a group of different phosphoproteins, mostly involved in protein binding and transport. In this dataset, we found proteins known to be involved in nitrogen response as differentially phosphorylated, including the high-affinity nitrate transporter NRT2.1^49,50^ and AMT1.3^69^. In our experimental conditions, site T464 in AMT1.3 was phosphorylated. Since phosphorylation of this site inhibits ammonium transport^69^, the increased phosphorylation status at this site may be related to the fact that our experimental conditions focused on nitrate transport in *Arabidopsis* roots. We also identified phosphoproteins associated with signaling pathways and transcription factors, uncovering regulatory networks linked to transcriptomic changes occurring later in the response to nitrate. The corresponding genes have not been characterized in the context of nitrate responses. For instance, one potentially novel regulatory factor is ABA-responsive element binding protein 3 (AREB 3), whose phosphoprotein levels are increased in response to nitrate treatments. This transcription factor is involved in ABA signaling^99,100^. Nitrate triggers ABA accumulation at the root tip^101^ and several studies indicate that ABA plays a role in regulation of lateral root growth by nitrate^102^.

NRT1.1/NPF6.3 is the main nitrate sensor described to date^31,39^. Interestingly, we observed important differences in the phosphoproteome of *chl1-5* mutant as compared to wild-type plants in response to nitrate treatments. These results emphasize that NRT1.1/NPF6.3, calcium and kinases/phosphatases make-up the canonical nitrate-signaling pathway of *Arabidopsis* roots. Recent evidence showed that NRT1.1/NPF6.3 and PLC activity are required for nitrate-induced increases in cytoplasmic Ca2+ levels^45^. Our results indicated that PLC2 phosphoprotein levels increased in response to nitrate. Nitrate signaling pathways also involve CBL–CIPK complex and CPK–NLP regulatory network (reviewed by ^48^).

The transcriptomic nitrate-response was not mirrored at phosphoproteomic levels. This lack of correlation between mRNA and protein or phosphoprotein levels has been documented in other nitrogen-phosphoproteomic studies^49,50^, plant responses to different stimulus^52,53^ and other organisms^103^. Furthermore, post-translational regulation does not require a change in gene expression or *de novo* protein synthesis. Post-translational control is faster, allowing rapid adaptation to environmental changes. Interestingly, the genes coding for nitrate-modulated phosphoproteins identified in this study are highly co-expressed across many different experimental conditions but not regulated by nitrate treatments^104^. This finding suggests that this group of genes is functionally related and regulated at the mRNA level in response to several endogenous or exogenous cues. In the context of nitrate responses, the products of these genes are regulated at the post-translational level, uncovering a new layer of control that enables signal crosstalk and fine tuning. Our results highlight the need for integrated analysis and data sets at different levels to decipher plant responses to environmental cues.

It is now clear that auxin plays a central role in the plant root response to changes in nitrate availability. Nitrate regulates primary root growth, lateral root initiation and elongation. Auxin in turn is key during root development^96^, particularly in initiation and growth of lateral roots^93^. Several reports show that auxin signaling, biosynthesis, transport, and accumulation are affected during nitrate responses^72,84^, and transcriptomic analyses demonstrate that genes involved in auxin response are controlled by nitrate^7,16,71^. The main nitrate transporter NRT1.1/NPF6.3 can also transport auxin^84,86^. A recent study also showed that NRT1.1/NPF6.3 negatively regulates the *TAR2* auxin-biosynthetic gene and the *LAX3* auxin-influx transport gene at low nitrate concentrations, repressing lateral root development^105^. These results suggest that an interplay between nitrate signaling and auxin transport occurs at different levels^106–108^. Consistent with these findings, we found that the molecular function “auxin transport” was overrepresented in our cluster and network analyses. We showed that dephosphorylation of PIN2 in a novel phosphosite is a part of a regulatory mechanism for RSA responses triggered by nitrate. The phosphorylation/dephosphorylation of PIN proteins at specific sites (serine or threonine) located in their higher loops has been shown to play important roles in modulating PIN functions^88,91,92,109^, including trafficking^110^. We showed that phosphorylation/dephosphorylation of PIN2 at S439 is important for PIN2 plasma membrane localization in epidermal and cortical cells in response to nitrate. Previous studies have shown that changes in PIN2 membrane localization and polarity interfere with PIN2 function in auxin transport with an impact on RSA during alkaline stress^111^ or low phosphate^112^. In addition, recent studies showed that kinase cascade modules 3′-PHOSPHOINOSITIDE-DEPENDENT PROTEIN KINASE 1 (PDK1)-D6 PROTEIN KINASEs (D6PK) and PDK1-AGC1 kinase PROTEIN KINASE ASSOCIATED WITH BRX (PAX) regulate auxin distribution through PIN phosphorylation^113,114^. The phosphorylation of PIN affects PIN-mediated auxin transport, which controls plant growth^113^ and other developmental processes^114^. These results suggest that PIN phosphorylation is part of a regulatory switch that strictly controls the directional transport of auxin and subsequent growth or developmental processes.

Our results suggest a model (Figure 9) in which nitrate promotes dephosphorylation of PIN2, which then impacts localization and auxin transport. Modulation of PIN2 function could affect growth of primary and lateral roots for optimal nutrient uptake. Beyond this new regulatory mechanism involving PIN2 protein, our phosphoproteomics results identify novel proteins, which may be interesting targets for future studies or biotechnological developments for improved nitrogen use-efficiency or crop yield.

**Figure 9.**
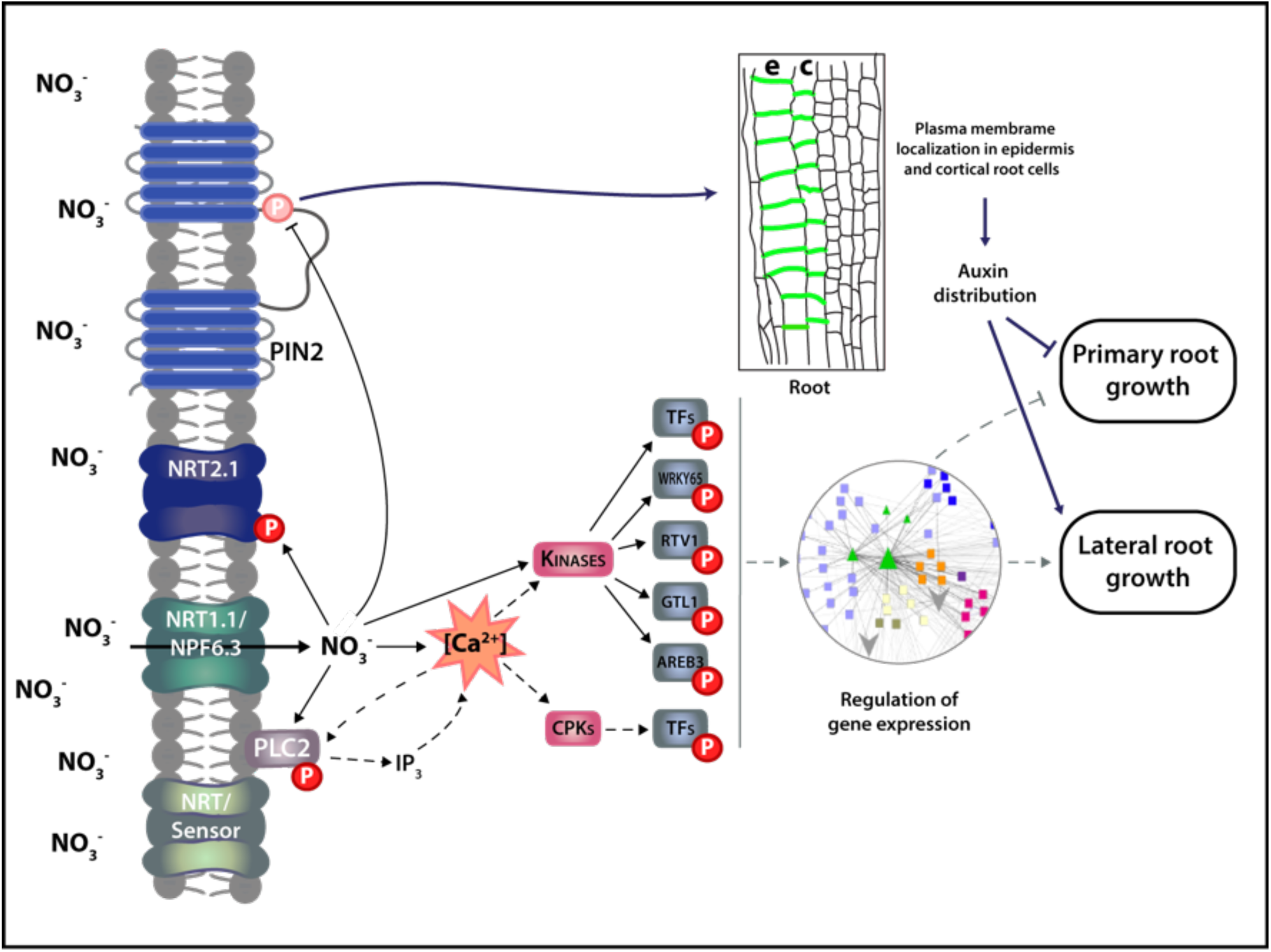
Schematic model of the role of PIN2 and phosphorylation in root nitrate responses. Nitrate treatments increase cytosolic calcium levels which interact with calcium binding proteins such as kinases in the nitrate signaling pathway. Kinases phosphorylate protein targets such as transcription factors that can mediate changes in gene expression in response to nitrate treatments. Nitrate can also cause dephosphorylation of specific proteins. We showed nitrate treatments promote PIN2 dephosphorylation at S439 which impact membrane localization and polarity of the PIN2 protein. This post-translational regulatory mechanism is important for modulation of primary root growth and lateral root density in response to nitrate treatments. Arrows and bar-headed solid lines represent activation or inhibition in response to nitrate, respectively. Dashed arrows indicate proposed connections.

## Acknowledgements

This work was supported by Millennium Institute for Integrative Biology—iBio (Iniciativa Científica Milenio-MINECON), Fondo de Desarrollo de Areas Prioritarias (FONDAP) Center for Genome Regulation (15090007), Fondo Nacional de Desarrollo Científico y Tecnológico (FONDECYT) 1141097 (to RAG) and 1171631 (to AV). We would like to thank Unidad de Microscopía Avanzada UC (UMA UC).

## Methods

### Plant material and growth conditions

*Arabidopsis thaliana (L*.*) Heynh. Columbia-0* accession plants were used as wild type genotype in all experiments. The transgenic lines *PIN2::PIN2*^*wt*^*-GFP, PIN2::PIN2*^*S439D*^*-GFP* and *PIN2::PIN2*^*S439A*^*-GFP* were introduced into *eir1-1* background (*pin2* null mutant plants), using strategies described previously. PIN2 lines were generated by Gybson assembly as described^115^ using pGREEN backbone vector^116^. *PIN2::PIN2*^*wt*^*-GFP* line was generated by insertion of *mGFP5* into *PIN2* coding sequence at nucleotide 1215 from ATG (between Thr405 and Arg406)^94,117^. For PIN2 promoter, 2178 bp upstream of the start codon was used. For the generations of *PIN2::PIN2*^*S439D*^*-GFP* and *PIN2::PIN2*^*S439A*^*-GFP* lines Serine 439 was replaced by aspartate (*PIN2*^*S439D*^) or alanine (*PIN2*^*S439A*^) by site-directed mutagenesis using Gibson Assembly.

Seeds were sterilized using 50% chlorine solution for 7 minutes and washed with sterile distilled water three times. Then, 1,500 *Arabidopsis* seedlings were placed into a hydroponic system (Phytatrays) with MS-modified basal salt media without N (Phytotechnology Laboratories, M531) supplemented with 1 mM ammonium as the only N source. Plants were grown under long-day photoperiods (16h light/ 8h dark and a temperature of 22 °C) for 14 days using a plant growth incubator (Percival Scientific, Inc.). At day 15, plants were treated with 5 mM KNO3 or 5 mM KCl for different time periods as indicated, using a protocol described previously^7^. For the phenotypic study of root response to nitrate, plants were grown as described above and were treated with 5 mM KNO3 or 5 mM KCl for 3 days.

### Root architecture analysis

For root phenotyping, plants were scanned in plates using an Epson Perfection V700 Photo scanner, and root length were measured using Fiji (v1.52). Initiating and emerging lateral roots were analyzed using DIC optics on a Nikon Eclipse 80i microscope, as described^7^. The data were statistically analyzed in the Graph Pad Prism 5 Program.

### Protein extraction, phosphopeptide enrichment and mass spectrometry analysis

Proteins were isolated from 1 g of frozen tissue per sample in each experimental condition (2-3 biological replicates). Sample preparation and protein extraction were performed using previously described methods^52,53^. Phosphopeptide enrichment was performed using 1% (w/v) colloidal CeO2 into an acidified peptide solution at 1:10 (w/w). Mass spectrometry (MS) and peptide identification were based on protocols described previously^60^. Briefly, the generated spectra were analyzed on LTQ Velos linear ion trap tandem mass spectrometer (Thermo Electron) and phosphorylation sites were identified into a specific amino acid within a peptide by using the variable modification localization score in Agilent Spectrum Mill software^118^. Proteins were grouped based on their shared, common phophopeptides using principles of parsimony to address redundancy in proteins. Proteins classified within the same group share the same subset of phosphopeptides. Phosphorotein levels were quantified using spectral counting, as described previously^52,53^. MS data were normalized using the total number of spectral counts (SPC) for each MS run. Expressed phosphoproteins were defined by at least one SPC, after the application of quality score cutoff in MS analysis, in minimum two of the tree biological replicates. To identify nitrate-regulated phosphoproteins in *Arabidopsis* roots, raw data were log transformed and quantile normalized using R studio (https://rstudio.com) and MEV (http://mev.tm4.org/) software. The statistical multifactor analysis of variance (significance: *p* < 0.05) and *pos-hoc* analysis (significance: *p* < 0.1) were performed using R-studio. Encoded genes for phosphoproteins showing a similar pattern were analyzed and visualized using the average-linkage hierarchical clustering performed in Cluster 2.11 software, as described^119^.

### Gene network analysis

*Arabidopsis* encoded genes from our phosphoproteomics data were used. The gene network was generated by integrating different information, including protein-protein interactions from BioGRID^74^, predicted protein-DNA interactions of *Arabidopsis* TFs (DapSeq)^75,76^, *Arabidopsis* metabolic pathways (KEGG), and miRNA-RNA, as described previously^16^. This analysis also included predicted regulatory connections between phosphoproteins that we detected and kinase families. A kinase-substrate analysis, identifying the most significant phosphorylation motifs and their predicted kinases were performed using the Motif-X algorithm^77^ and PhosPhAt Kinase-Target interactions database^78,120^. The resulting network was visualized using CYTOSCAPE software^79^. Cluster and Gene Ontology analysis into the network were achieved with ClusterMaker^121^ and Bingo^121^ tools for biological networks in cytoscape, respectively.

### Phos-Tag PAGE and western immunoblotting

Affinity-based SDS PAGE identification of phosphorylated PIN2 isoforms were performed based on the protocols by Komis et al.^122^ and the protocol given by the Phos-tag manufacturer (FUJIFILM Waco Chemicals) with slight modifications. *Arabidopsis* seedlings were grown on modified MS plates with ammonium as the only N source for 7 days. Next, seedlings (n≥40) were transferred to 5 mM nitrate amended agar plates and were incubated for 6 hours in light. Root samples were collected and homogenized in liquid N2 and then extracted with extraction buffer – 50 mM Tris-HCl, pH 7.5, 150 mM NaCl, 0.5% Triton X-100, 10 µM MG-132 and 0.1 mM PMSF - supplemented with protease and phosphatase inhibitor cocktails (Roche). Buffer volumes were adjusted to fresh weight (100 µL/100 mg tissues). Homogenized samples were centrifuged (4°C, 15 min, 14000 rpm) and the supernatant was aliquoted (50µg protein/20-25 µL) and incubated at 45°C for 5 min in the presence of SDS loading buffer. Next, samples were loaded onto an acrylamide, Bis-Tris/HCl gel containing 50 µmol/L Phos-tagTM (AAL-107) pendant and Zn^2+^ as cation. Electrophoresis were run at 15 mA/gel for 5∼ 6 hours or until the proteins are nicely separated (usually until the 25 kDa prestained protein marker just exit the gel assembly). Next gels were incubated for 30 minutes in transfer buffer containing 10 mM EDTA and were blotted to PVDF membranes using Tris/Glycine transfer buffer (25 mM Tris, 192 mM Glycine, 5% methanol). After blocking with 5% milk in TBST, the membrane was probed with α-PIN2 antibody (1:1000) for 2 h (RT) followed by α-rabbit IgG-HRP (1:15000) (Amersham) for 1 h (RT). After treating the membrane with Supersignal West Femto western chemiluminescent HRP substrate (Thermo Fisher Scientific), luminescent signals were detected using a liquid nitrogen–cooled charge-coupled device camera (Biorad). Digital images were analyzed, and signals were quantified using the Fiji software.

### Confocal microscopy and image analysis

*eir1-1* mutants complemented with *PIN2::PIN2*^*wt*^*-GFP, PIN2::PIN2*^*S439A*^*-GFP, PIN2::PIN2*^*S439D*^*-GFP* were grown in modified MS plates with 1 mM ammonium as the only N source. 2 week-old plants were treated with 5 mM nitrate or without treatment (control conditions) for 2 hours. Roots were stained with propidium iodide (PI) and mounted on a slice for microscopic analysis. Images were acquired with Zeiss LSM880 confocal microscope with Airyscan equipped with a 40×Plan-Apochromat water immersion objective. Fluorescence signals for GFP (excitation 488 nm, emission 507 nm) and PI (excitation 536 nm, emission 617 nm) were detected. Microscopy

For image quantification (PIN2-GFP fluorescence intensity measurements) maximum intensity projections of confocal pictures were used in epidermis and cortex cells. Images were handled and analyzed with Zeiss blue (v3.1) and Fiji (v1.52) software. The experiment was performed with 3 biological replicates and 3-4 roots per experimental conditions were analyzed in each replicate.

---

**Figure S1.**
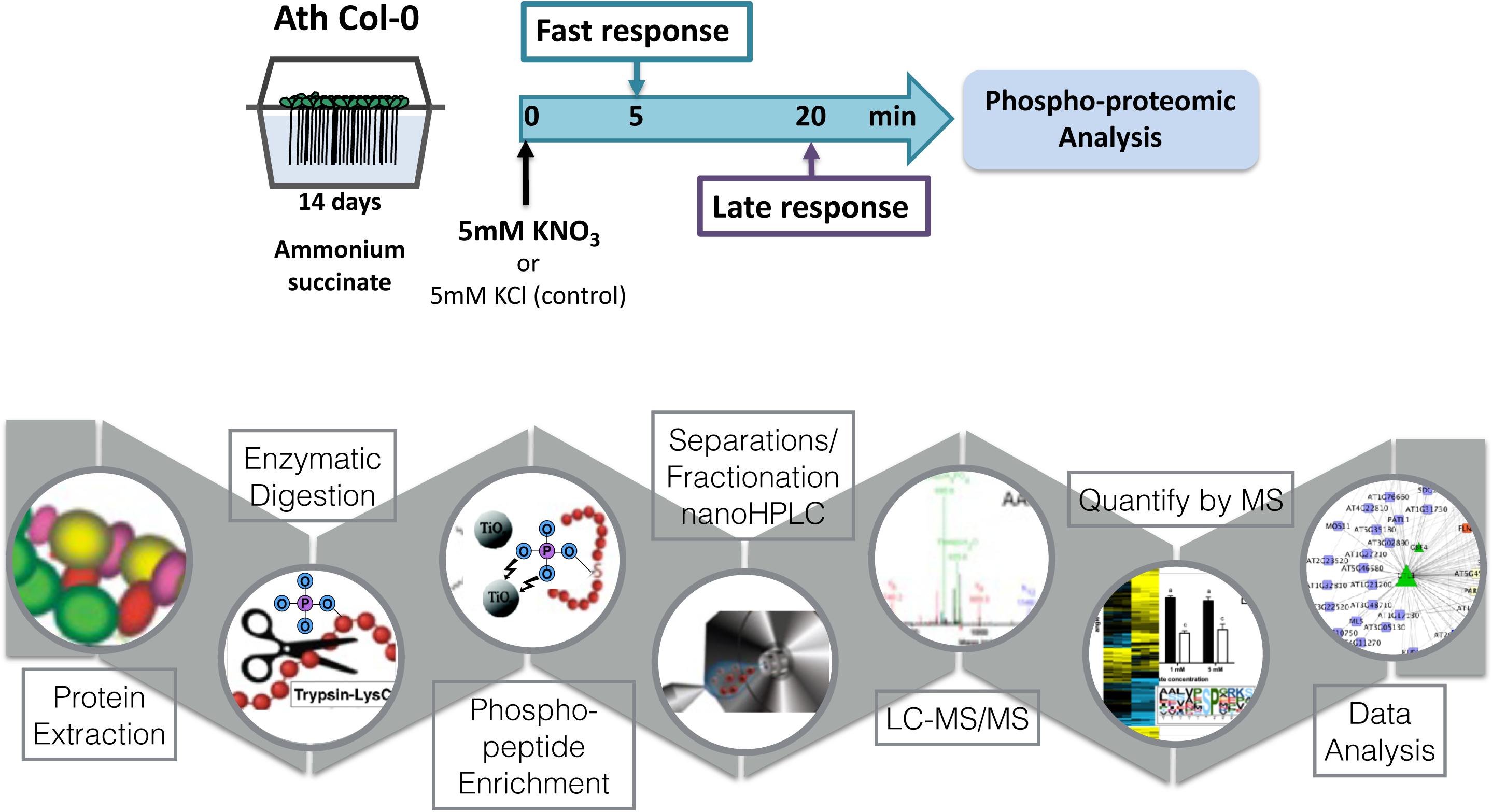
An overview of the root phosphoproteomic experiment. (A) Experimental design to identify changes in phosphoproteins in response to nitrate treatments in *Arabidopsis* thaliana roots. (B) Phosphoproteomic strategy for the enrichment, quantification and analysis of phosphopeptides.

**Figure S2.**
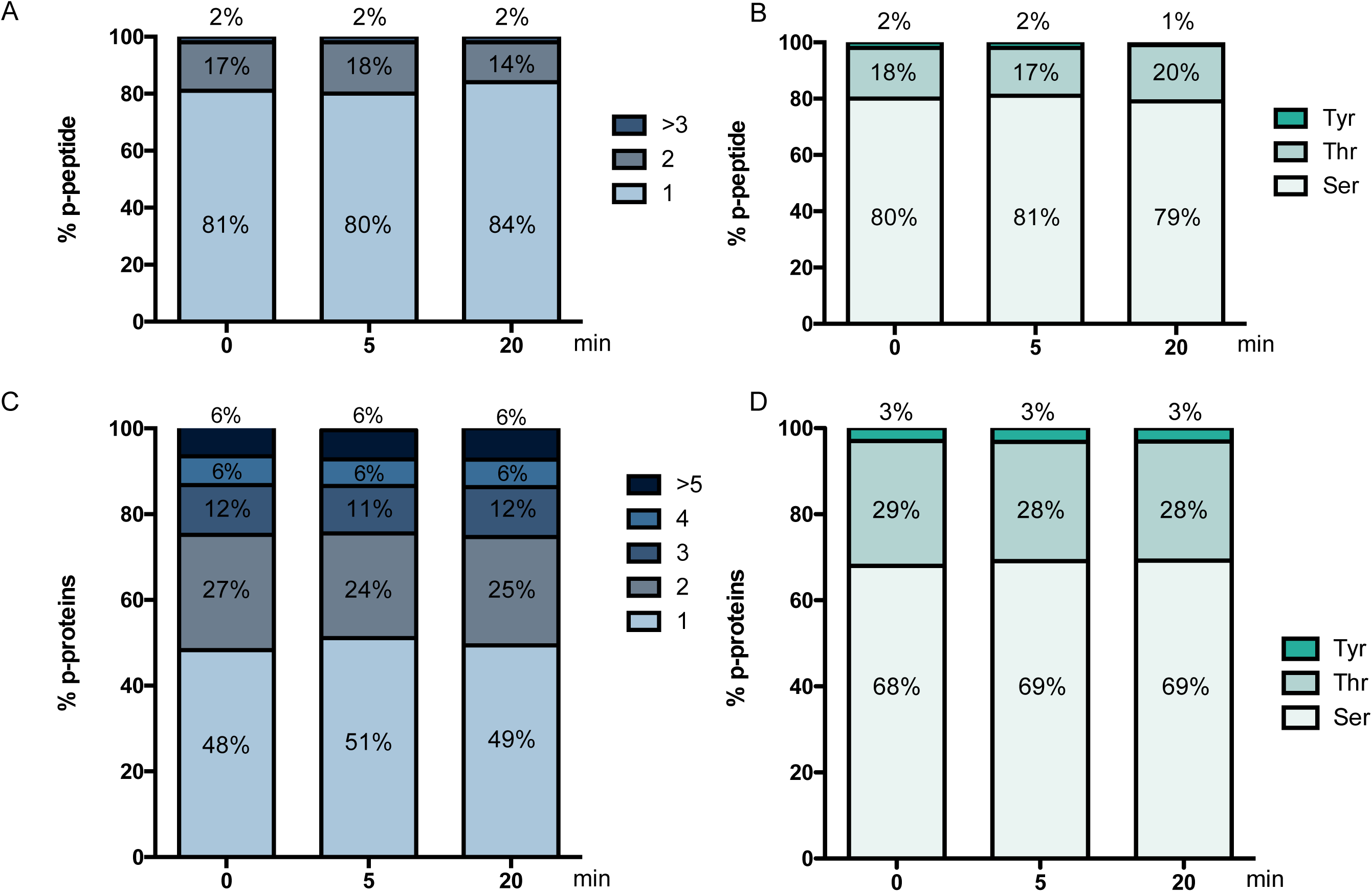
An overview of the root phosphoproteomic analysis. (A) Distribution of the number of phosphosites per peptide for the indicated experimental data set (0, 5 or 20 minutes after nitrate treatment). (B) Relative distribution of phosphorylated residues in each peptide for indicated experimental data set. (C) Distribution of the number of phospho-peptides per protein for indicated experimental data set. (D) Percentage of phospho-proteins that present each phosphorylated residues for indicated data set.

**Figure S3.**
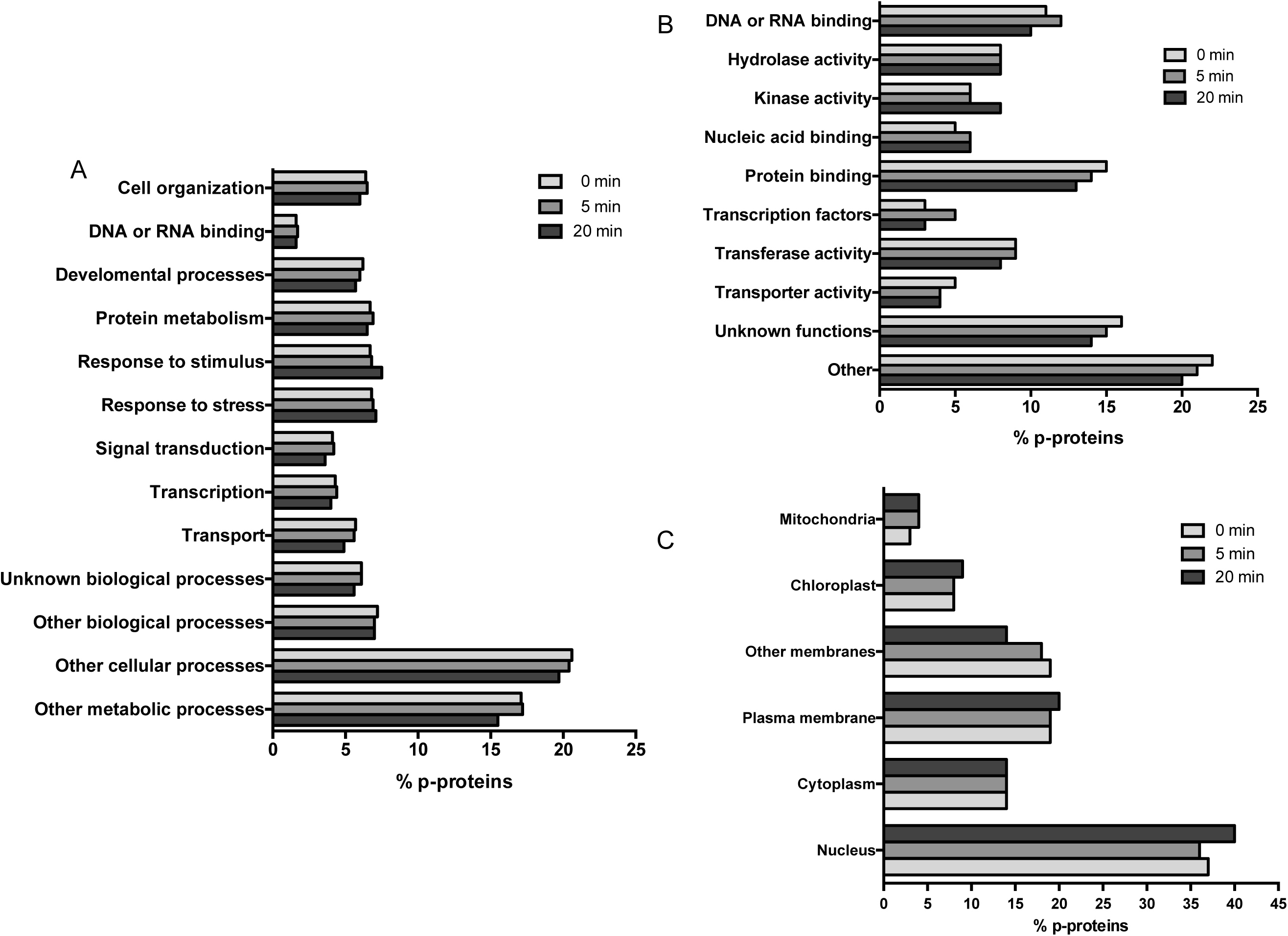
Distribution of phosphoproteins across GO categories: biological process (A), cellular functions (B) and sub cellular compartment (C) for each experimental data set.

**Figure S4.**
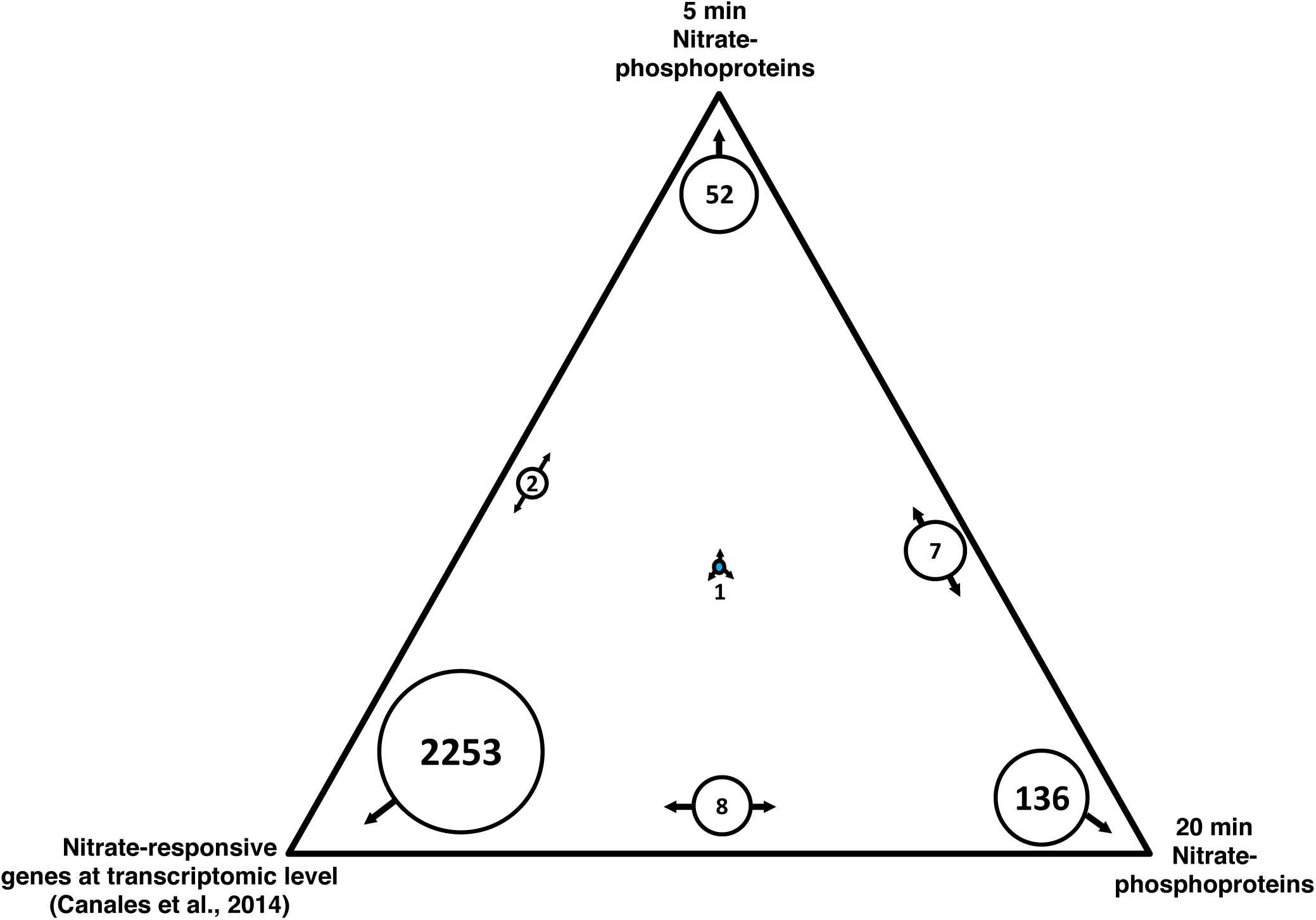
Comparison between phosphorproteomics and transcriptomic dataset in response to nitrate. The list of encoded genes in response to nitrate at transcriptomic (data-set 27 affymetrix experiment Canales et al., 2014) and phosphoproteomic (our work) levels were represented using the SUNGEAR tool (Poultney et al., 2007). The triangle shows each data set at the vertices: transcriptomic data-set, phosphoproteomic data (5 min), phosphoproteomic data (20 min).The size of the circles inside the figure are proportional with the number of genes in each data set, as indicated by the arrows around the circle. The number of genes is indicated in each circle.

**Figure S5.**
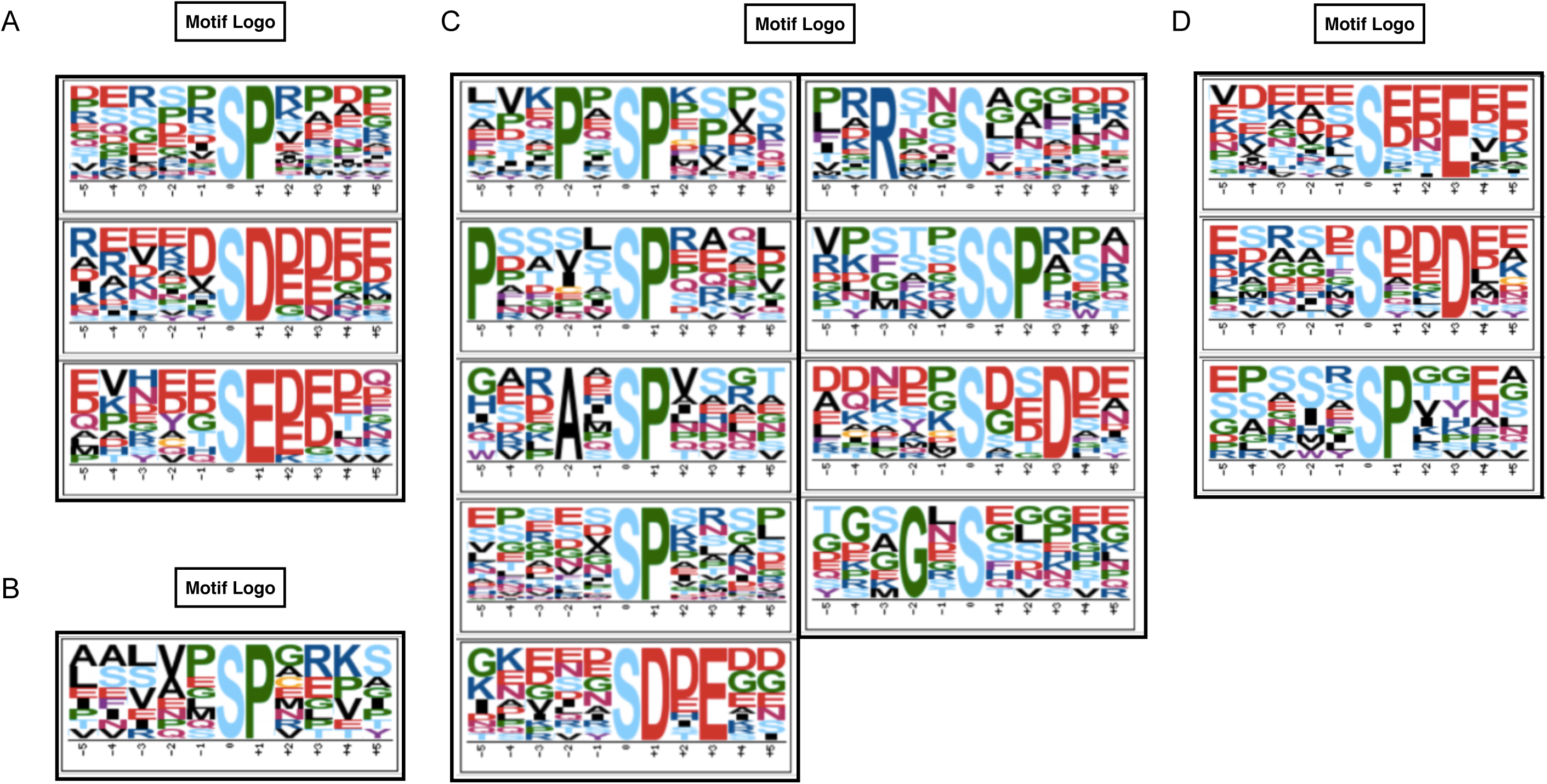
Motif-X analysis of nitrate-responsive phosphopeptides. Amino acid sequences from -5 to +5 residues (the phosphorylated residue was 0) were scanned with the Motif-X algorithm. Motif list of phosphopeptides that met the criteria for phosphoproteins up-regulated (A) and down-regulated (B) by nitrate at 5 min. Motif list of phosphopeptides that met the criteria for phosphoproteins up-regulated (C) and down-regulated (D) by nitrate at 20 min See table S1 for the complete set of nitrate-regulated phosphoproteins.

**Figure S6.**
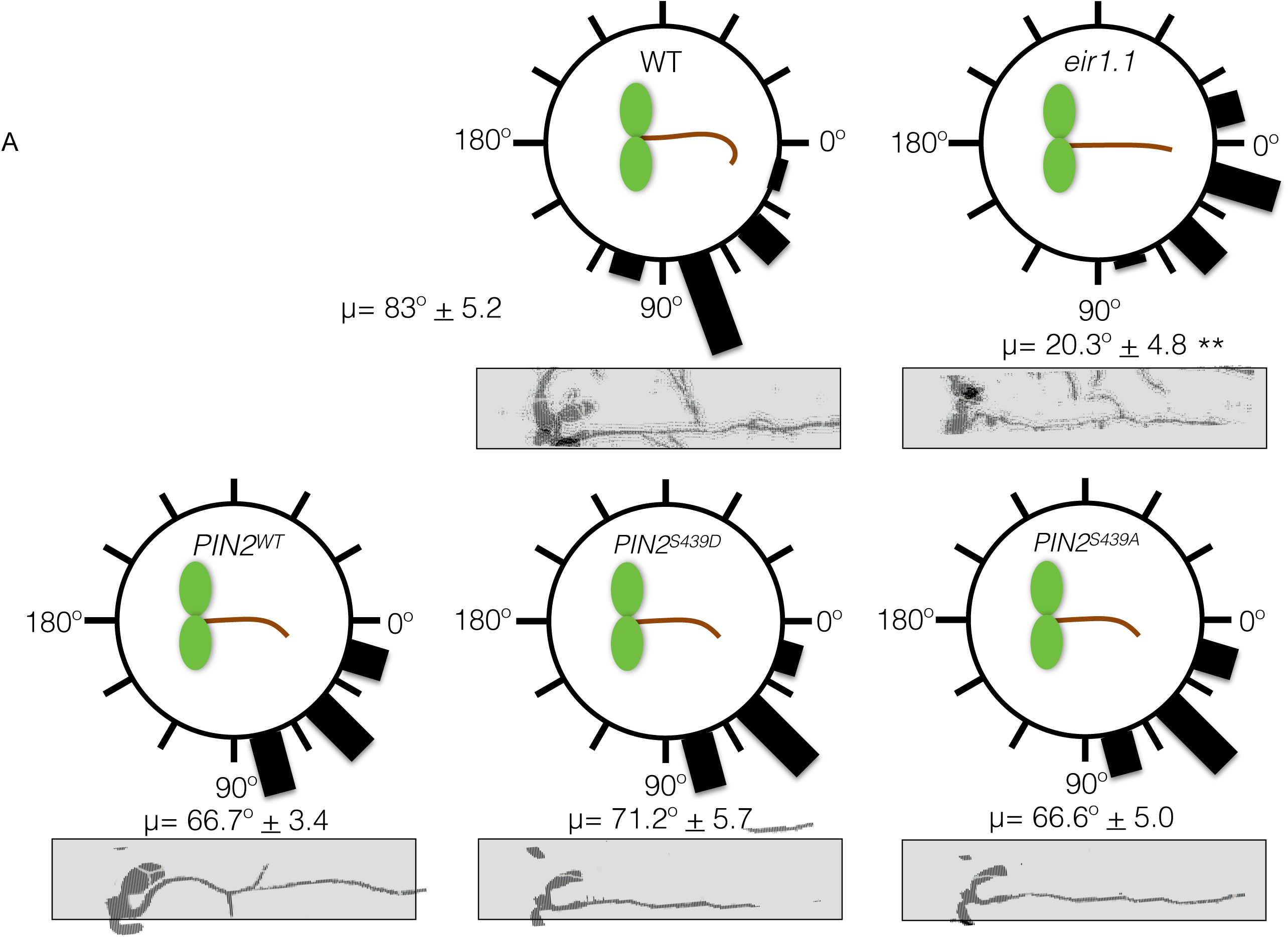
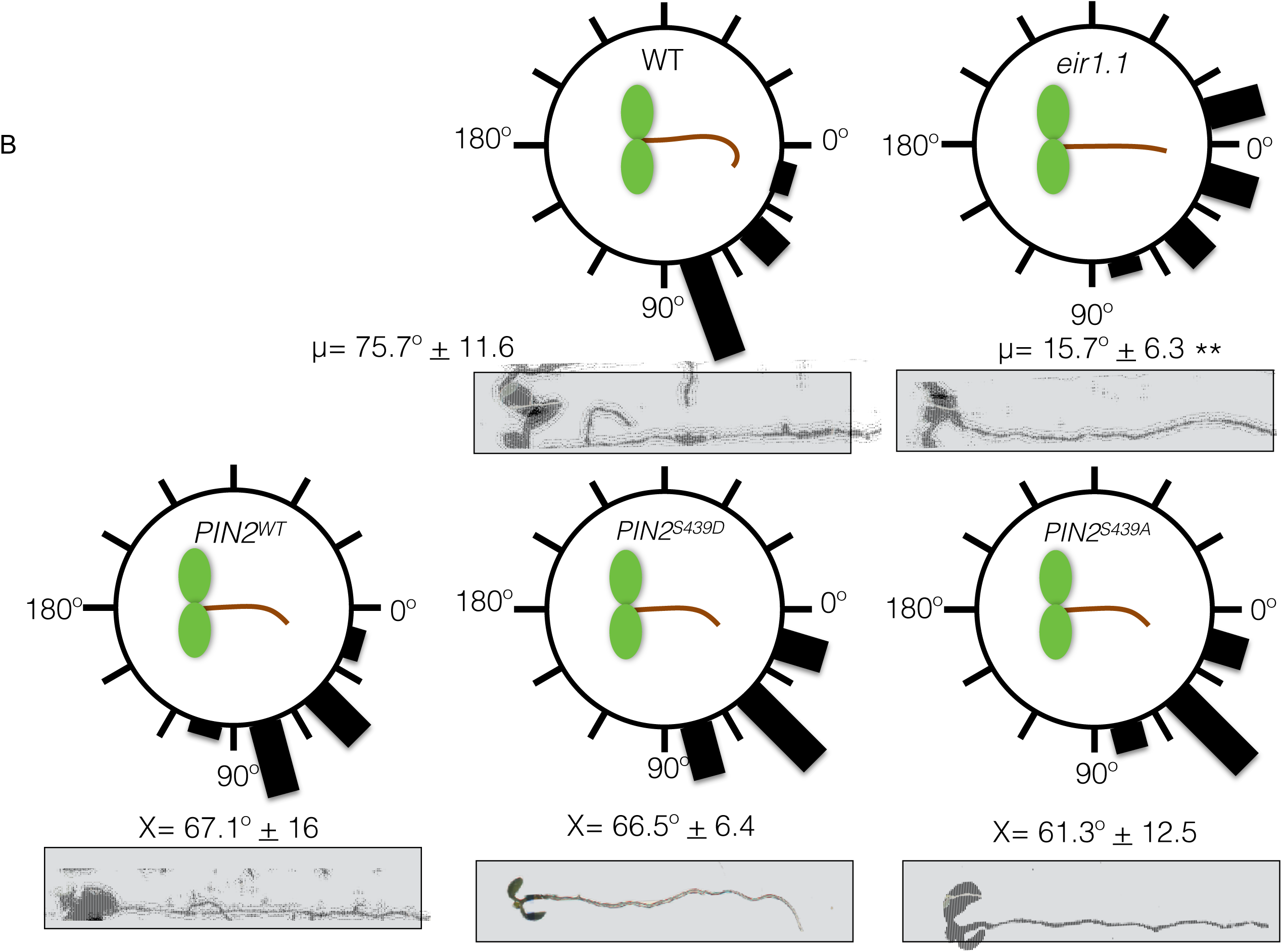
Phosphoprotein PIN2 (S439) in root gravitropic response. pin2 null mutant background (*eir1-1)* complemented with non-modified or wild type version of PIN2 (PIN2^wt^), phospho-null (S439A) or phospho-mimic (S439D) versions of PIN2 (PIN2^S439A^ or PIN2^S439D^, respectively) were grown in MS agar plate supplemented with 0 mM (A) or 5 mM (B) nitrate. At day 7, plates were rotated 90° and root curvature was measured at 24 hours with imageJ software. The figure shown a histogram of root gravitropic response where the length of black bars indicates the relative frequency of plants with the corresponding classes of angle. The average (µ) plus standard deviation of seedlings scored per line is indicated below each circle. Asterisk indicate statistically significant difference between means (\*\**p* < 0.01).

